# Phenotypic assessment and genetic validation of *Plasmodium falciparum* molecular markers associated with malaria chemoprevention in Senegal

**DOI:** 10.1101/2025.11.05.686777

**Authors:** Katelyn Vendrely Brenneman, Wesley Wong, Mamy Yaye Die Ndiaye, Stephen Schaffner, Bassirou Ngom, Karina Bellavia, Imran Ullah, Amy Gaye, Djiby Sow, Mame Fama Ndiaye, Mariama Toure, Nogaye Gadiaga, Aita Sene, Awa Bineta Deme, Baba Dieye, Mamadou Samb Yade, Khadim Diongue, Younouss Diedhiou, Jules François Gomis, Mouhamadou Ndiaye, Mamadou Alpha Diallo, Ibrahima Mbaye Ndiaye, Bronwyn MacInnis, Dyann F. Wirth, Daouda Ndiaye, Sarah K. Volkman

**Affiliations:** Harvard T.H. Chan School of Public Health, Boston, MA, USA; International Research & Training Center in Applied Genomics and Health Surveillance (CIGASS), Cheikh Anta Diop University, Dakar, Senegal; The Broad Institute, Cambridge, MA, USA; Department of Parasitology, Faculty of Medicine and Pharmacy, Cheikh Anta Diop University, Dakar, Senegal; Senegal National Malaria Control Program; Simmons University, School of Nursing

## Abstract

Drug resistance in *Plasmodium falciparum* threatens to undermine malaria control and elimination efforts. Senegal is a malaria-endemic country that has implemented successive antimalarial and chemopreventive drug-based strategies for two decades. Sulfadoxine-pyrimethamine (SP) is used for chemoprevention in Senegal for intermittent preventive treatment in pregnancy (since 2004) and SP plus amodiaquine (AQ) is used for seasonal malaria chemoprevention (SMC, since 2013). Using whole genome sequence (WGS) data from malaria patient samples from health facilities across Senegal (2006 – 2022), we observed near fixation of *Pfdhfr* triple mutant and fluctuation in *Pfdhps* and *Pfcrt* mutation frequencies over time. It is unclear how these mutations influence drug resistance and fitness phenotypes in natural isolates; therefore, we evaluated natural parasite isolates with different *Pfcrt, Pfmdr1, Pfdhps*, and *Pfdhfr* haplotypes. Parasites were culture-adapted and phenotyped for antimalarial drug susceptibility and competitive growth (fitness).

*Pfcrt* CVIET + A220S + Q271E + N326S + R371I and *Pfcrt* CVIET + A220S + Q271E + I356T + R371I mutants were significantly more resistant to monodesethyl-amodiaquine (md-AQ) compared to *Pfcrt* wild-type (WT) and *Pfcrt* CVIET + A220S + Q271E + R371I mutants. *Pfdhfr* triple mutants were significantly more pyrimethamine (PYR) resistant than *Pfdhfr* WT and revealed a range of phenotypes, but this was not explained by *Pfgch1* copy-number. *Pfdhps* A437G parasites were significantly more sulfadoxine (SDX) resistant compared to *Pfdhps* wild-type and *Pfdhps* S436A mutants, suggesting that A437G is a key mutation for SDX resistance. Competitive growth assays between *Pfdhfr-Pfdhps* mutants revealed that *Pfdhps* mutations do not always result in fitness costs. Ongoing phenotypic assessment and genetic validation of these mutations in a Senegalese background is necessary to assess the impact of drug pressure, identify evolving genetic determinants of drug resistance, and provide molecular markers for ongoing surveillance to monitor and guide the use of drug-based interventions.

**AUTHOR SUMMARY:** Drug resistance is a major concern for both preventing and treating malaria, especially in Africa where most malaria cases and deaths occur. Since 2013, Senegal has been giving children under 10 years old a combination of sulfadoxine-pyrimethamine plus amodiaquine to prevent malaria during the transmission season, called Seasonal Malaria Chemoprevention (SMC), and plans to continue expanding its use. However, there is evidence from genetic surveillance that drug resistance mutations are present in Senegal which could render this antimalarial drug combination ineffective. Here we use natural *P. falciparum* isolates obtained from Senegalese patients that represent the extant parasite population to evaluate the consequences of evolving mutations on antimalarial drug resistance and fitness phenotypes. This study is one of the first to use natural parasites to assess the impact of naturally derived mutations on drug resistance and fitness phenotypes. Our results provide evidence that certain combinations of drug resistance mutations impact both parasite drug resistance and fitness, and therefore need to be closely monitored and can inform optimal antimalarial combinations for the prevention or treatment of malaria. This work informs the ongoing evolution of resistance and fitness phenotypes in malaria endemic settings that are introducing new multi first line therapies (MFTs) and SMC interventions that have been used for decades in Senegal. Our approach creates a framework for using genetic surveillance data to form a hypothesis, which can then be phenotypically tested by measuring the resistance and fitness levels of genetically diverse natural parasite isolates.

## INTRODUCTION

Malaria remains a global health concern, with the majority of malaria cases and deaths occurring in African children under 5 years old (1). Antimalarial drugs continue to be one of the main methods to combat *Plasmodium falciparum* malaria; they are used to both treat malaria infections and prevent malaria infection. Many different antimalarial drugs have been used throughout history, and drug resistance has emerged to every antimalarial drug in use (2). Genetic surveillance of drug resistance mutations can identify resistant parasites and determine whether resistance has emerged locally or if resistance mutations have spread from other regions (3,4). By tracking drug resistance mutations and identifying their sources, genetic surveillance is a critical tool for monitoring and predicting the risk of drug resistance, controlling the spread of drug resistance, and determining whether drug-based interventions will be effective in certain regions.

Typically, genetic surveillance is conducted using genotyping data of known resistance markers and therefore relies on knowledge of the molecular basis of drug resistance for antimalarial drugs of interest. For several antimalarials, the basis of drug resistance has been well established by previous studies: mutations in *P. falciparum* chloroquine resistance transporter (*Pfcrt*) are associated with chloroquine (CQ) resistance (5,6). The molecular basis of sulfadoxine (SDX) and pyrimethamine (PYR) drug resistance is associated with accumulations of point mutations in the genes encoding the targets of SDX and PYR, dihydropteroate synthase (*Pfdhps*) and dihydrofolate reductase (*Pfdhfr*), respectively (7,8) that encode essential enzymes involved in the folate biosynthesis pathway required for DNA synthesis. However, there are several antimalarials with unknown or partial molecular markers of resistance, including quinolines such as amodiaquine (AQ) and lumefantrine (LUM); both likely involve mutations in *Pfcrt* and the multi-drug resistance *Pfmdr1* gene and other unidentified mutations for resistance (9–12). Or in the case of artemisinin, there is a major marker of resistance, *Pfkelch13* (13), but there are likely other mutations that confer resistance (14–17). Since the genetic determinants of several antimalarial drugs have not been fully elucidated and may involve multiple genes or compensatory mutations (18), genetic surveillance of known drug resistance markers can only provide an estimate of drug resistance in the population, highlighting the need to further characterize the drug response and fitness of parasites that have evolved under drug pressure and carry different combinations of known mutations.

Senegal is a malaria-endemic country that has implemented successive antimalarial drug-based strategies for treatment and prevention and has conducted genetic surveillance for several decades. CQ was used as a frontline antimalarial until 2003, when it was replaced by sulfadoxine-pyrimethamine (SP) plus AQ. SP + AQ remained the frontline antimalarial combination until Senegal transitioned to Artemisinin Combination Therapy (ACT) in 2006. ACT use began with artesunate and amodiaquine (AS-AQ) in 2006, followed by artemether-lumefantrine (AL) in 2008; both ACTs remain the current frontline antimalarials. For chemoprevention, Senegal currently uses SP for intermittent preventative treatment in pregnancy (IPTp, since 2004), and uses SP + AQ for Seasonal Malaria Chemoprevention (SMC) for children (since 2013). SMC is given by community health workers in 3-5 monthly rounds (depending on local malaria incidence and the duration of transmission season) for all children under 10 years of age (19).

For the past two-decades in Senegal, genetic surveillance has been used to assess how antimalarial drug usage for treatment and prevention has affected parasite populations. The frequency of drug resistance markers of CQ (mutations in *Pfcrt*), artemisinin (mutations in *Pfkelch13*), AQ (mutations in *Pfmdr1* and *Pfcrt*), and SP (mutations in *Pfdhfr* and *Pfdhps*) were examined for trends of increasing drug resistance that could signal a reduction in antimalarial drug efficacy. Molecular surveillance revealed drastic changes over time; the *Pfcrt* K76T mutation declined from 76% to 26% prevalence after the withdrawal of CQ in 2003 but since 2014 has rebounded to 49%.

The frequency of *Pfdhfr* mutations (N51I, C59R, S108N) have been steadily increasing from 42% since 2000 and are nearing fixation (95%). The frequency of *Pfdhps* A437G has been fluctuating from 17% to 72% over the last 10 years, while *Pfmdr1* (N86, Y184F, D1246) appears to have stabilized at a frequency of 56% (19). These mutation frequencies from genetic surveillance strongly suggest that parasites in Senegal are rapidly evolving in response to antimalarial drug use. Because both drug resistance and parasite fitness can impact treatment efficacy and influence treatment strategies, it is important to understand the effects of known drug resistance mutations on these phenotypes in natural parasite isolates. Therefore, we performed “phenotypic surveillance”, assaying natural parasite isolates *in vitro* to determine their drug resistance and fitness phenotypes.

Natural parasite isolates are representative of parasite genomes that are currently circulating in the field and can be culture-adapted for *in vitro* phenotyping. Therefore, natural parasite isolates are extremely useful for estimating the extent of drug resistance in natural populations and for identifying mutations that are driving drug resistance and fitness phenotypes. *In vitro* drug susceptibility assays are one approach to assess parasite drug responses. While they have limitations, they provide extremely valuable information about how certain mutations or haplotypes affect parasite drug response. A previous study assessing the *in vitro* drug susceptibility of 45 culture-adapted parasite isolates to 12 different antimalarials showed that these isolates exhibit a variety of drug resistance phenotypes. This study also confirmed the roles of *Pfcrt, Pfdhfr,* and *Pfmdr1* as mediators of resistance, but also found several novel signals of PYR resistance-associated selection on chromosome 6 and 12 (20). However, since this 2012 study, there have been many changes to drug pressure in the population and SDX or SP were not included in that analysis; consequently, the role of *Pfdhps* and its interactions with known and novel resistance mutations was not investigated.

SMC was first deployed across the Sahel subregion of Africa in 2012 for children aged 3-59 months, and as of 2023, the use of SMC has been expanded to 19 African countries that have highly seasonal malaria transmission (1). In addition, restrictions on the number of monthly cycles or age were removed so that SMC could be given to children at high risk of severe malaria during peak malaria transmission season (21). Given the expanded use of SMC (both geographically and in adding additional monthly cycles), especially in areas of Africa where drug resistance is present, and the significant increase in the prevalence of *Pfdhfr* triple mutants only two years after the implementation of SMC in several African countries (22), genetic surveillance has increased in an effort to monitor known drug resistance markers. However, translating this genetic surveillance data into actionable knowledge that can be used by public health professionals who implement policy changes based on this data remains a challenge. Therefore, it is important to understand how the effectiveness of the continued use and expansion of SMC and other interventions that use SP (IPTp and perennial malaria chemoprevention (PMC)) could be compromised by *Pfdhfr* and *Pfdhps* mutations.

In this study, we used parasites that were collected from patients in Senegal and whole genome sequenced over the past two decades to identify major haplotypes of known drug resistance markers present in the population (19,23). Based on these results, we culture-adapted and phenotyped parasite isolates with certain combinations of these mutations that were all present in the natural population, including *Pfcrt* (codon positions 74-76, A220S, Q271E, N326S, I356T, and R371I), *Pfdhfr* (N51I, C59R, S108N), *Pfdhps* (S436A, A437G, A613S), and *Pfmdr1* (N86Y, Y184F, D1246Y) to determine what role these alleles were playing in parasite fitness or drug resistance. We found that *Pfcrt* CVIET A220S + Q271E + N326S + R371I and *Pfcrt* CVIET + A220S + Q271E + I356T + R371I mutants were significantly more resistant to md-AQ compared to wild-type *Pfcrt* parasites. We saw a range of SDX resistance phenotypes regardless of *Pfdhps* genotype, but the key mutation for sulfadoxine resistance appeared to be *Pfdhps* A437G. There was not a clear correlation between fitness and drug resistance, but our findings provide evidence that certain combinations of drug resistance mutations are impacting both parasite drug resistance and fitness and should be closely monitored.

## RESULTS

### Analysis of drug resistance haplotypes and selection of isolates representing the overall population haplotypic diversity

*Plasmodium falciparum* samples were collected between 2006 and 2022 from febrile patients from six sampling locations within Senegal: Pikine, Thiès, Diourbel, Kaolack, Kolda, and Kédougou. These samples were whole genome sequenced and samples that were monogenomic (single genome infection) were used for further analysis (1879 samples). To represent the genetic diversity of Senegal parasites as a whole, we selected monogenomic natural parasite isolates for culture-adaptation collected from two sites: Thiès (low transmission area; 2021 reported annual incidence was 2.8 cases per 1000) and Kédougou (high transmission area; 2021 reported annual incidence was 536.5 cases per 1000) (24). Parasites were chosen for culture-adaptation based on their combination of alleles at *Pfcrt* (codon positions 74-76, A220S, Q271E, N326S, I356T, and R371I), *Pfmdr1* (N86Y, Y184F, D1246Y), *Pfdhfr* (N51I, C59R, S108N), and *Pfdhps* (I431V, S436A, A437G, K540E, A581G, A613S), which defined their combined *Pfcrt*, *Pfmdr1*, *Pfdhfr*, and *Pfdhps* haplotype. Based on these 19 different genomic sites of the combined haplotype, there was a total of 524,288 unique haplotypes (2^19^) that could possibly exist, however, we only found 239 unique haplotypes in our dataset, likely because these haplotypes are linked and not random (e.g., *Pfcrt* M74I, N75E, and K76T are commonly found together), some combinations are incompatible, and many genomic sites are nearing fixation in Senegal (e.g., *Pfdhfr* N51I, C59R, S108N). Out of the 239 haplotypes found in our dataset, 208 haplotypes had less than 10 parasites with that haplotype (134 haplotypes had only 1 parasite). Therefore, we selected 33 parasites for culture-adaptation that represented 26 haplotypes (10.9% of all haplotypes in our dataset); these 33 parasites represented 63.9% of all parasites in our dataset (supplemental table 1, supplemental figure 1).

### Determining Pfmdr1 and Pfgch1 copy-number variation

Once culture-adapted, the 33 parasites were whole genome sequenced again to confirm their sequences; all parasite samples were confirmed to have the same genomic regions of interest pre-culture adaptation and post-culture adaptation. Parasites were also assayed for copy-number variations (CNVs) in *Pfgch1* and *Pfmdr1* (table 1). While no parasites showed any CNVs in *Pfmdr1* (associated with mefloquine resistance), some parasites had some copy-number amplifications in *Pfgch1*, which encodes a GTP cyclohydrolase that is the first and rate-limiting enzyme in the folate synthesis pathway (*Pfdhfr* and *Pfdhps* are key enzymes in the later stages of this pathway). *Pfgch1* has been shown to have extensive copy-number polymorphisms with some amplifications of *Pfgch1* occurring only in the gene and some amplifications only occurring in the promoter region (25–27); both types of amplifications are likely a result of selection by antifolate drugs SDX and PYR acting in different geographical regions (28,29). Therefore, we assayed parasites for amplifications in the *Pfgch1* gene and the *Pfgch1* promoter. We found that 3 parasites had promoter amplifications (all had triple promoter amplifications, supplemental figure 2), and 1 parasite had a *Pfgch1* gene amplification (Th052.16). Studies have shown that increased copy-numbers of *Pfgch1* have a variable effect on PYR EC_50_ values and could also be a compensatory mechanism for the fitness cost imposed by *Pfdhfr* and *Pfdhps* (28,30,31). Further functional characterization is needed to determine whether amplifications in the *Pfgch1* promoter or the *Pfgch1* gene result in an increase in *Pfgch1* expression.

**Table 1:**
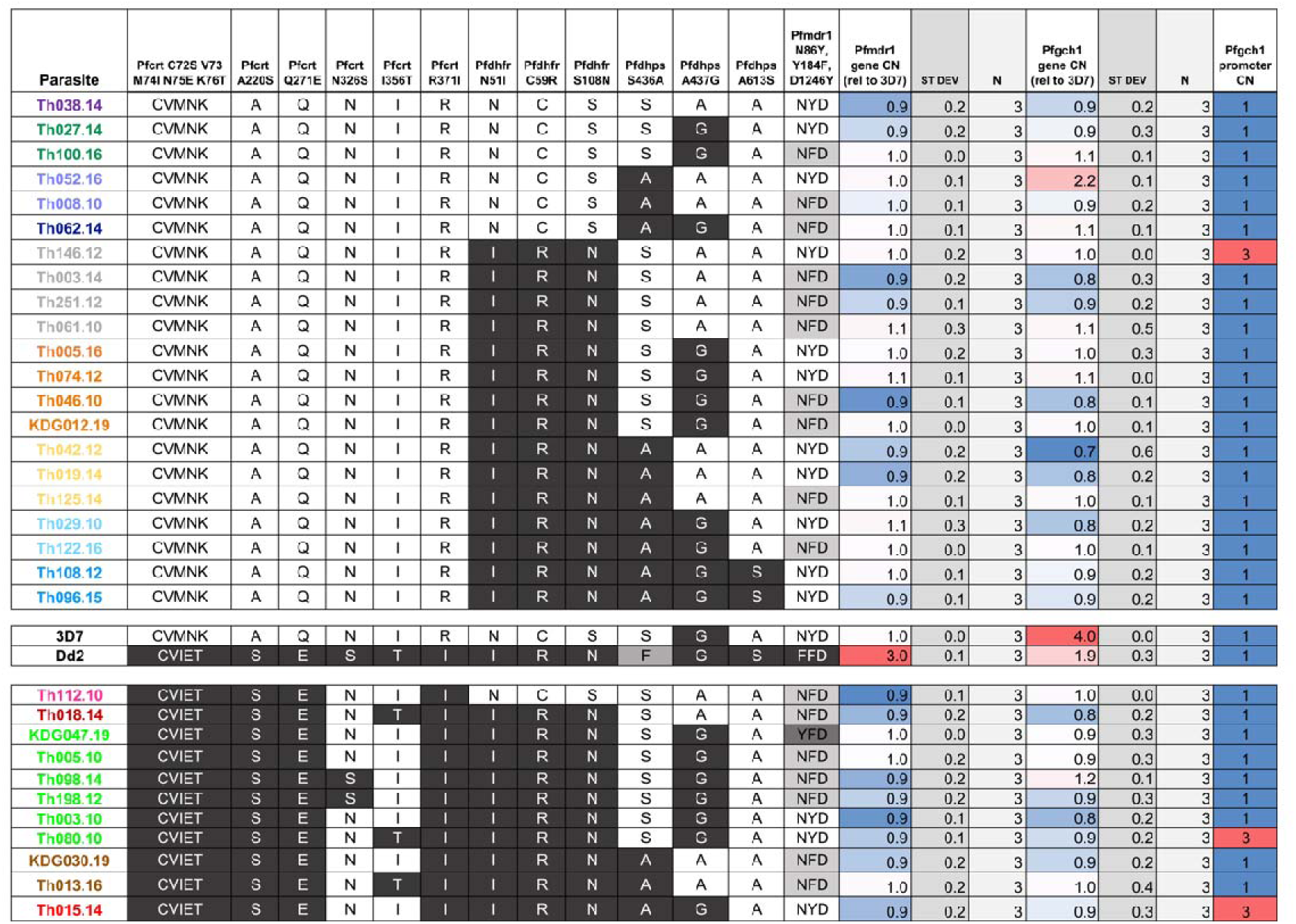
Genotypes and CN variations for 33 natural parasite isolates from Senegal and 2 reference lines selected for phenotypic assessment.

### Parasite drug susceptibility phenotyping

To determine whether the different *Pfcrt*, *Pfmdr1*, *Pfdhfr*, and *Pfdhps* combined haplotypes have an impact on drug resistance, we investigated the drug susceptibility of our culture adapted parasite isolates to a panel of antimalarial drugs that have been used for antimalarial therapy or chemoprevention in Senegal. The set of culture-adapted parasites (and 3D7 and Dd2 reference strains) were phenotyped for *in vitro* drug susceptibility to chloroquine (CQ), mefloquine (MQ), lumefantrine (LUM), piperaquine (PIP), dihydroartemisinin (DHA), amodiaquine (AQ), quinine (QN), monodesethyl-amodiaquine (md-AQ), pyrimethamine (PYR), sulfadoxine (SDX), and sulfadoxine-pyrimethamine (SP). No statistically significant difference in EC_50_ values was seen amongst parasites for MQ, LUM, PIP, DHA, AQ, and QN, however, we did see statistically significant differences in EC_50_ values for CQ, md-AQ, PYR, SDX, and SP (supplemental table 2 and supplemental figure 3).

### The role of Pfcrt mutations in conferring chloroquine and amodiaquine resistance

One of the most well-studied relationships between drug resistance and causative mutation is CQ and *Pfcrt* K76T. However, Senegal has not used CQ since 2003 and yet genetic surveillance data revealed an increase in *Pfcrt* K76T allele frequency beginning in 2014 (19). It is also well documented that there is a fitness cost to parasites carrying *Pfcrt* K76T mutations (32–34), so why are *Pfcrt* K76T allele frequencies rising in Senegal? We hypothesize that AQ resistance (which likely involves mutations in *Pfcrt*, *Pfmdr1*, and other unidentified mutations for resistance) could be driving this change. AQ is a component of both an ACT frontline antimalarial treatment (AS-AQ) and is also a component of SMC (SP + AQ) and has been extensively used throughout Senegal for the past decade; however, according to national policy, AS-AQ and SMC are not used in the same regions, AL is used as the frontline antimalarial instead. Importantly, artemisinin resistance has not yet appeared in Senegal (35). To better understand the drug susceptibility profile of parasites with the most common *Pfcrt* + *Pfmdr1* haplotypes present in our Senegal dataset, we exposed parasites to CQ and md-AQ (the active metabolite of AQ) to determine their EC_50_ values. Unsurprisingly, we found that mutant *Pfcrt* CVIET parasites had significantly higher CQ EC_50_ values compared to *Pfcrt* wild-type CVMNK parasites; parasites with the *Pfcrt* CVIET + A220S + Q271E + I356T + R371I (Cam783) haplotype had the highest CQ EC_50_ values compared to all other haplotypes (figure 1A), as was shown in a previous study (32). *Pfcrt* CVIET parasites with or without *Pfmdr1* mutations did not have significantly different CQ EC_50_ values, suggesting that *Pfmdr1* mutations do not play a role in CQ resistance in this set of parasites.

**Figure 1.**
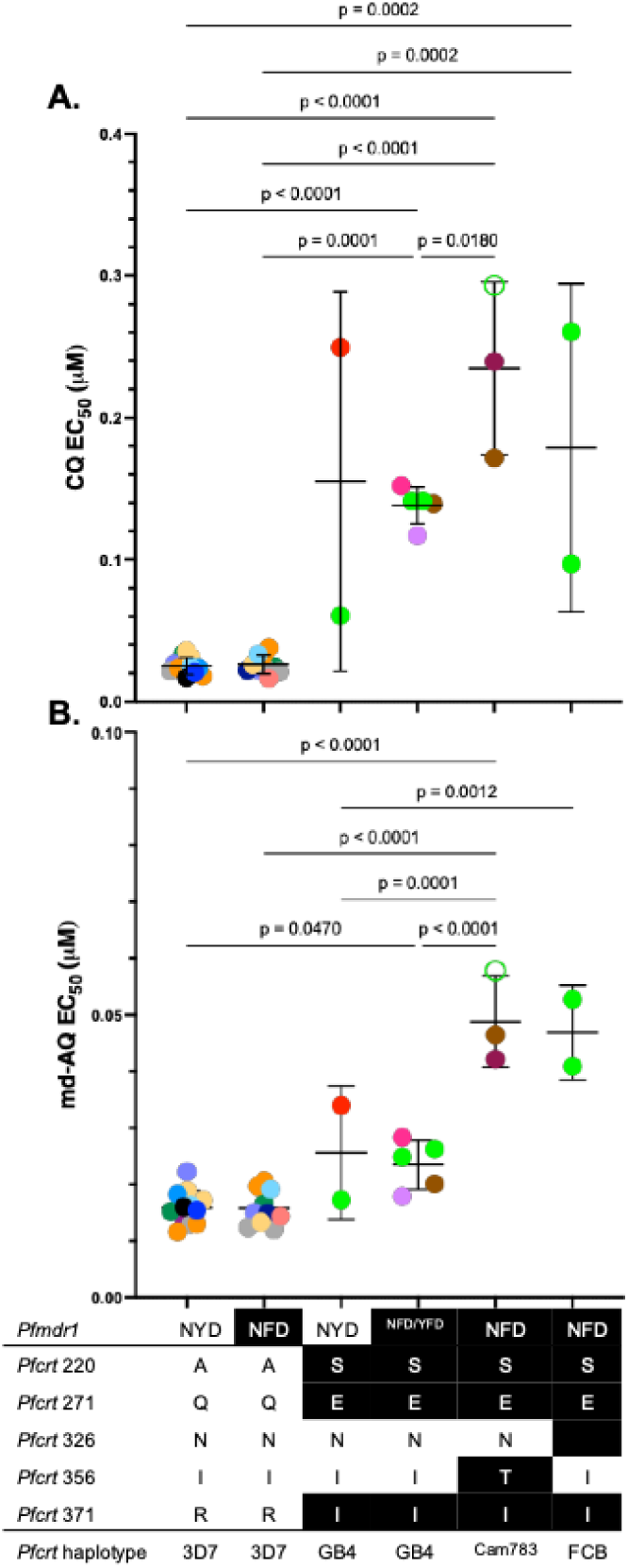
A. Chloroquine (CQ) EC_50_ comparisons and B. monodesethyl-amodiaquine (md-AQ) EC_50_ comparisons. (A) Parasites with *Pfcrt* CVIET mutations (GB4, Cam738, and FCB haplotypes), regardless of *Pfmdr1* mutations, are statistically significantly more resistant to CQ. (B) Parasites with *Pfcrt* CVIET mutations (GB4, Cam738, and FCB haplotypes), regardless of *Pfmdr1* mutations, are statistically significantly more resistant to md-AQ; GB4 parasites were 1.3-fold more md-AQ resistant than 3D7 and Cam783 parasites were 3.3-fold and FCB parasites were 3.1-fold more md-AQ resistant than 3D7. Parasite with an open green circle is *Pfmdr1* NYD.

We found that mutant *Pfcrt* CVIET parasites had increased md-AQ EC_50_ values compared to *Pfcrt* wild-type CVMNK parasites (figure 1B). Interestingly, we found that *Pfcrt* CVIET A220S + Q271E + I356T + R371I (Cam783 haplotype, p=0.0001) and *Pfcrt* CVIET A220S + Q271E + N326S + R371I (FCB haplotype, p=0.001) parasites had significantly higher md-AQ EC_50_ values compared to CVIET + A220S + Q271E + R371I (GB4 haplotype), which suggests that the *Pfcrt* N326S and I356T mutations could be playing a key role in mediating AQ susceptibility. However, none of the parasite isolates tested had elevated EC_50_ values that would be considered AQ resistant (∼0.06 μM (36)). Several studies have shown that *Pfcrt* mutations, especially the mutant *Pfcrt* CVIET haplotype, could be one of the potential drivers of AQ resistance (37–41), but it is most likely that AQ resistance is driven by multiple mutations. There is also evidence that in addition to *Pfcrt* mutations, *Pfmdr1* mutations and copy-number variations modulate drug resistance (10,37,42). Notably, none of the parasites in our dataset have *Pfmdr1* copy-number amplifications or *Pfmdr1* N1042D or D1246Y mutations (table 1), which means we cannot determine their contribution to drug resistance. However, in our dataset, parasites with *Pfmdr1* N86Y or Y184F mutations did not have significantly greater md-AQ EC_50_ values compared to *Pfmdr1* wild-type parasites (either with or without *Pfcrt* CVIET). Therefore, these specific *Pfmdr1* mutations (N86Y or Y184F) do not seem to explain the elevated md-AQ EC_50_ values. However, there are likely other undiscovered contributors to AQ resistance; *Pfcrt* and *Pfmdr1* mutations play a role and should be closely monitored, but alone do not predict AQ resistance (43). Based on these results, *Pfcrt* mutant parasites that include either N326S or I356T mutations should be closely monitored in Senegal for potentially indicating decreased AQ susceptibility. Interestingly, parasites with *Pfcrt* N326S or I356T mutations are also more resistant to CQ compared to wild-type parasites at these loci (supplemental figure 4).

### *Pfcrt* allele population frequency over time

Given the decreased md-AQ susceptibility for parasites with *Pfcrt* N326S or I356T mutations, we wanted to determine the frequency of these mutations in Senegal over time. To do this, we investigated the full haplotypes of *Pfcrt* (amino acid codon positions 72-76, 220, 271, 326, 356, and 371) and *Pfmdr1* (amino acid positions 86, 184, and 1246) from 2006-2022 across all sites in Senegal and found that there were four major haplotypes in the 2160 samples: wild-type parasites (*Pfmdr1* NYD + *Pfcrt* 3D7, black), *Pfmdr1* mutant parasites (NFD + 3D7, gray), mutant *Pfcrt* parasites (NYD + Cam783, maroon), and mutant *Pfcrt* and *Pfmdr1* parasites (NFD + Cam783, red) (figure 2). Comparing the *Pfmdr1* and *Pfcrt* combined haplotypes over time for Senegal and for just Thiès and just Kédougou revealed similar trends over time (supplemental figure 5). The most common haplotype found throughout Senegal in 2022 was mutant *Pfmdr1* (NFD + 3D7) (21.8% [18.0, 25.6]), which we found to have no resistance to CQ or md-AQ *in vitro*. However, of the parasites that had *Pfcrt* mutations, the most common haplotype was mutant *Pfcrt* and *Pfmdr1* (NFD + Cam783, red) (15.7% [12.4,19.0]), which has been increasing in frequency since 2012 (2.4% [0.0,7.0]) and we found to be the least susceptible to CQ and md-AQ *in vitro*. The *Pfcrt* FCB (purple) haplotype had a similarly elevated md-AQ EC_50_ *in vitro*, however, its frequency throughout Senegal has been declining since 2014 (13.3% [3.4,2.3] in 2014 to 1.9% [0.01,0.3] in 2022). The *Pfcrt* Cam783 haplotype (red), along with the GB4 haplotype (green) have been reported in previous studies to be two of the most common African mutant *Pfcrt* haplotypes; these haplotypes were found to be susceptible to AQ and had minimal fitness costs compared to other mutant *Pfcrt* haplotypes (32,44). Based on our analysis of 10,037 samples from the MalariaGEN Pf7 dataset, the *Pfcrt* Cam783 haplotype is the most common mutant *Pfcrt* haplotype in West Africa, the GB4 haplotype is the most common mutant *Pfcrt* haplotype in Central and Eastern Africa, and the FCB haplotype is the most common in Northeastern Africa (supplemental figure 6).

**Figure 2.**
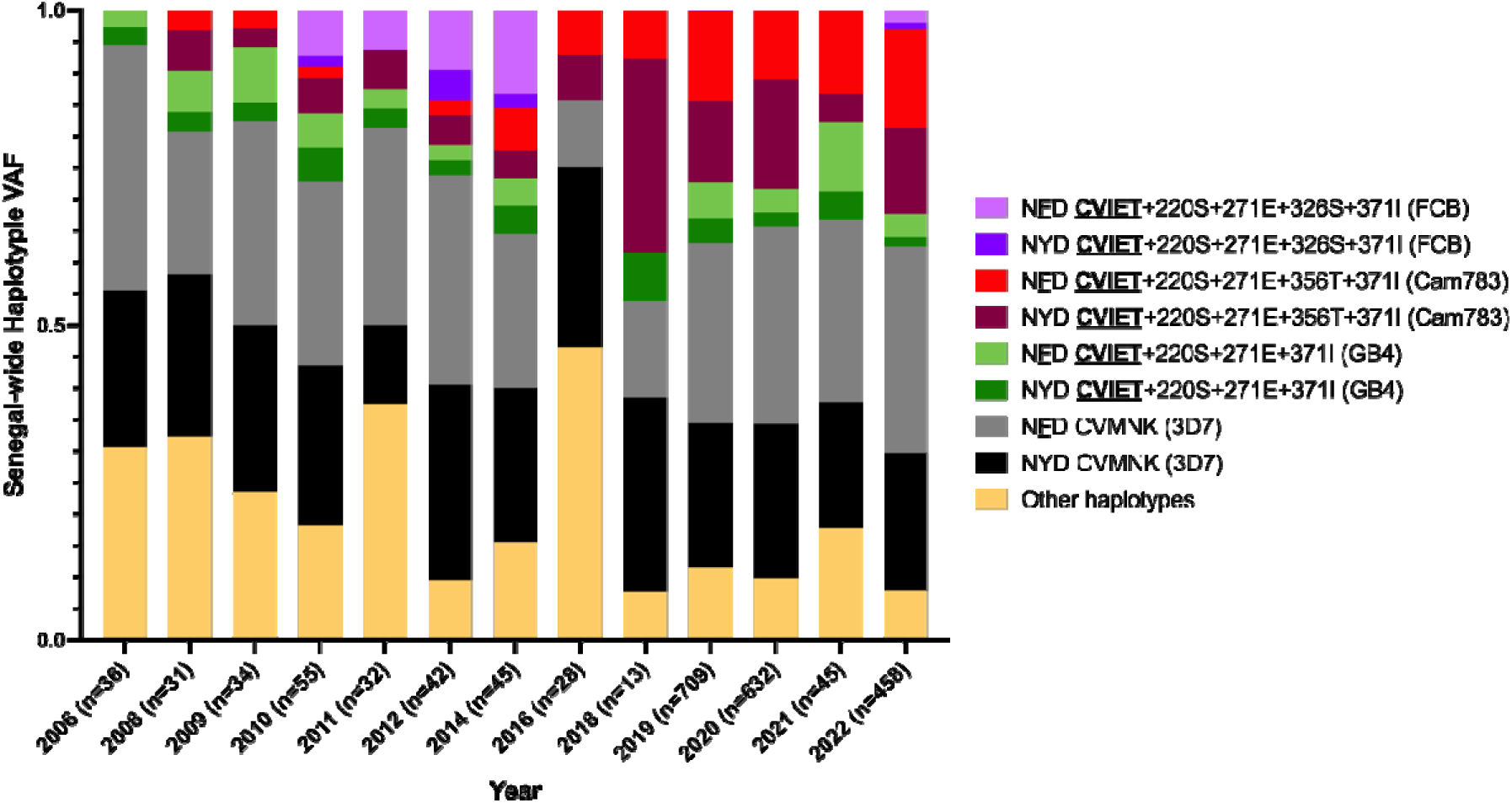
Haplotype variant allele frequencies (VAF) for *Pfcrt* and *Pfmdr1*. Samples collected from patients throughout Senegal were whole genome sequenced, 2160 monogenomic infections with genotype calls at *Pfcrt* and *Pfmdr1* were included in this dataset. The most common *Pfcrt* haplotype since 2006 is wild-type *Pfcrt* (3D7, gray/black), while the most common mutant *Pfcrt* haplotype is Cam783 (red). *Pfmdr1* NYD and NFD haplotypes remained even over time, but the NFD haplotype has a slightly higher frequency (0.47 [0.44,0.51]) than NYD (0.44 [0.41,0.48]) across all years throughout Senegal. The full list of haplotypes that are included in the “other haplotypes” category can be found in supplemental table 3.

### The in vitro fitness of Pfcrt alleles in the absence of drug

Next, we wanted to determine whether certain combinations of *Pfmdr1-Pfcrt* mutations had a fitness cost in these natural isolates. We determined that *Pfcrt* mutations have a large fitness cost in pairwise competition growth assays with *Pfcrt* wild-type parasites, therefore, we chose 11 *Pfcrt* mutant CVIET parasites that have various combinations of *Pfcrt* and *Pfmdr1* mutations to compete in pairwise competitions. These 11 parasites were each co-grown with another parasite in all unique combinations, resulting in 55 competitions. Parasites were set up 1:1 and co-grown until one parasite outcompeted the other, or after 30 days if there was no winner, the competition was called a tie.

Based on the competitive outcomes, parasites were ranked from most fit (i.e., won every competition and have a perfect winning percentage of 100%) to the least fit (i.e., lost every competition and have a winning percentage of 0%). Th080.10 (*Pfcrt* Cam783 + *Pfmdr1* NYD) was the most fit, while KDG047.19 (*Pfcrt* GB4 + *Pfmdr1* YFD) was the least fit. Perhaps unsurprisingly, a *Pfmdr1* NYD wild-type parasites was the most fit parasite; the fitness cost of *Pfmdr1* mutations has been previously shown (45). However, there was no fitness difference between NYD and NFD parasites; parasites with each genotype displayed a range of fitness phenotypes (figure 3A). Parasites from each of the different *Pfcrt* CVIET categories displayed a range of relative fitness (i.e., *Pfmdr1* NFD or YFD + GB4 had the least fit parasite and the third-most fit parasite). Cam783 and FCB haplotypes had the top two most fit parasites, suggesting that in the particular genetic background of those two parasites, N326S and I356T do not confer a fitness cost (figure 3B).

**Figure 3.**
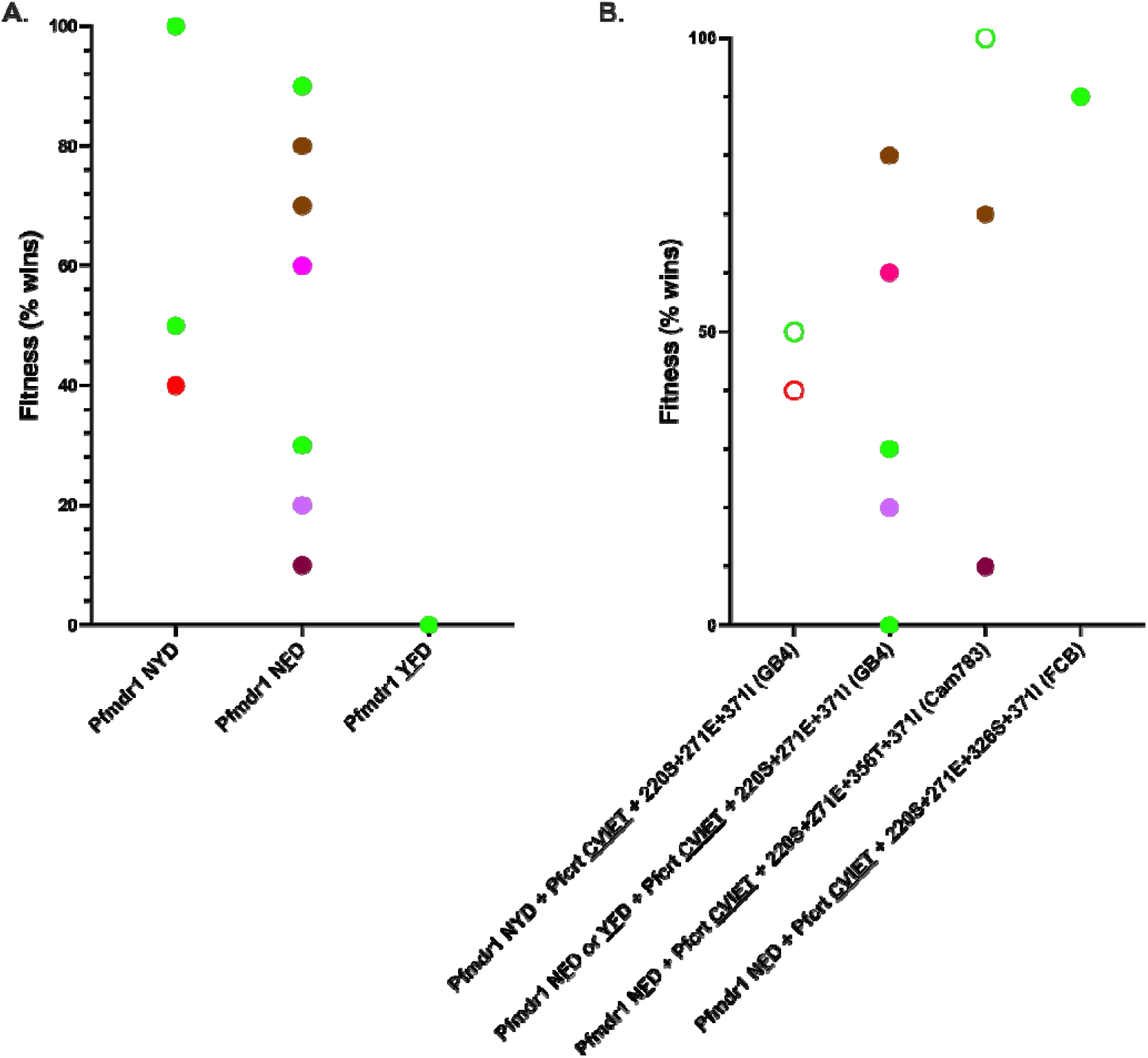
Relative fitness of 11 *Pfcrt* CVIET parasites based on A. *Pfmdr1* haplotype and B. *Pfcrt* and *Pfmdr1* haplotype. (A) Parasites with YFD mutations were the least fit compared to *Pfmdr1* wild-type (NYD) and *Pfdmr1* NFD parasites. (B) Amongst parasites with *Pfcrt* CVIET mutations, an I356T mutant parasite and a N326S mutant parasite were the two most fit. *Pfmdr1* NYD parasites are indicated by open circles.

### Allelic determinants of pyrimethamine and sulfadoxine resistance in Senegalese parasites

Parasites need folate to provide cofactors for many processes, including DNA synthesis (30,46). Parasites can salvage exogenous folate from the host and can also synthesize folate *de novo* (endogenous folate), however the efficiency of folate salvage varies between parasite strains (47). *Pfdhps*, the target of SDX, is an enzyme that catalyzes the conversion of endogenous folate precursors to dihydropteroate, a key intermediate in folate biosynthesis, which is then converted to tetrahydrofolate by *Pfdhfr*, the target of PYR. *Pfdhfr* can also convert exogenous folate precursors to tetrahydrofolate. The ability of *Pfdhfr* to convert both exogenous and endogenous folates, and therefore an ability to bypass the *Pfdhps* step, is likely why *Pfdhfr* mutations occur before *Pfdhps* mutations. Mutations in *Pfdhfr* accumulate in a step-wise fashion (S108N is often first, then N51I or C59R) and result in PYR resistance (48,49). Mutations in *Pfdhps* are selected for once parasites have at least two mutations in *Pfdhfr* and include I431V, S436A/F, A437G, K540E, A581G, and A613S/T. It is well known that the efficacy of SP is affected by mutations in *Pfdhfr* and *Pfdhps*, however the impact of various combinations of these mutations has not been characterized.

#### Pyrimethamine resistance

Our previous molecular surveillance study showed that the *Pfdhfr* triple mutant (N51I, C59R, S108N) is nearly fixed in Senegal (19). *Pfdhfr* is a well-studied and known driver of PYR resistance, and the accumulations of mutations in *Pfdhfr* are often the first steps in the evolution of SP resistance. To determine the impact of these *Pfdhfr* haplotypes on PYR resistance, we adapted parasites to para-aminobezoic acid (PABA)-free and folate-free media and phenotyped parasites with and without *Pfdhfr* triple mutations (N51I, C59R, S108N) and with and without combinations of *Pfdhps* mutations. In our data, we saw clearly elevated PYR EC_50_ values when comparing *Pfdhfr* **<u>IRN</u>** mutant parasites to *Pfdhfr* wild-type parasites (figure 4A). Interestingly, there is a range of PYR EC_50_ phenotypes amongst the *Pfdhfr* **<u>IRN</u>** mutant parasites, which suggests that other mutations or copy-number variations could also be driving these different PYR phenotypes.

**Figure 4.**
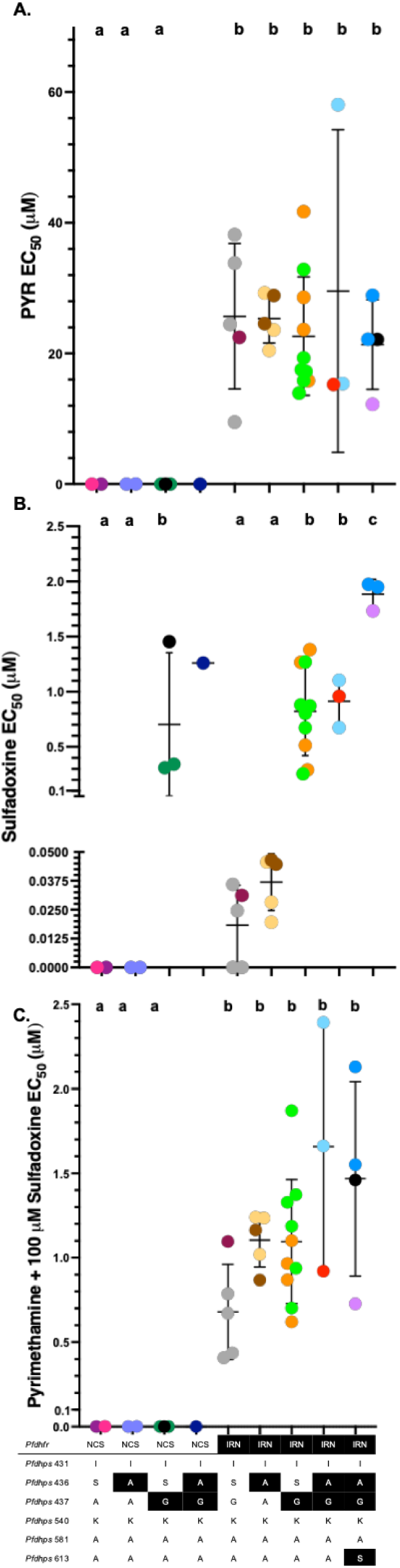
A. Pyrimethamine (PYR) EC_50_ comparisons, B. sulfadoxine (SDX) EC_50_ comparisons, and C. PYR + 100 μM SDX EC_50_ comparisons. (A) Parasites with *Pfdhfr* **<u>IRN</u>** mutations are statistically significantly more resistant to PYR compared to wild-type parasites. (B) Parasites with *Pfdhps* A437**<u>G</u>** mutations and S436**<u>A</u>** + A437**<u>G</u>** mutations are significantly more resistant to SDX compared to wild-type parasites and parasites with *Pfdhps* S436**<u>A</u>** mutations, while parasites with *Pfdhps* S436**<u>A</u>** + A437**<u>G</u>**+ A613**<u>S</u>** mutations are significantly more resistant to SDX compared to all other parasite *Pfdhps* haplotypes. (C) As parasites accumulate mutations in *Pfdhfr* and *Pfdhps*, overall, they become less susceptible to SP. Parasites with *Pfdhfr* **<u>IRN</u>** + *Pfdhps* S436**<u>A</u>** + A437**<u>G</u>** mutations are significantly more resistant to SP compared to all other parasite haplotypes. Letters indicate significant differences between in EC_50_ means between groups (p<0.05); groups labeled with the same letter (a, b, or c) are not significantly different from each other, while groups with different letters from each other are significantly different.

#### Sulfadoxine resistance

Given the near fixation of the *Pfdhfr* **<u>IRN</u>** triple mutation in Senegal and the continued use of SP, we next examined the effect of mutations in *Pfdhps*, a known driver of SDX resistance. Our set of parasites included several different *Pfdhps* haplotypes: *Pfdhps* wild-type (ISAKAA), *Pfdhps* S436A mutant (I**<u>A</u>**AKAA), *Pfdhps* A437G mutant (IS**<u>G</u>**KAA), *Pfdhps* S436A + A437G mutant (I**<u>AG</u>**KAA), and *Pfdhps* S436A + A437G + A613S mutant (I**<u>AG</u>**KA**<u>S</u>**). While these haplotypes have been reported before in West Africa (22,50), the impact of these mutations on drug resistance remains unknown. Parasites were adapted to PABA-free and folate-free media and assayed for SDX susceptibility (figure 4B). Parasites with the *Pfdhps* I**<u>A</u>**AKAA haplotype had the same SDX EC_50_ values as *Pfdhps* wild-type parasites. However, parasites with the *Pfdhps* IS**<u>G</u>**KAA and I**<u>AG</u>**KAA haplotype had significantly higher SDX EC_50_ values compared to wild-type and I**<u>A</u>**AKAA parasites (p<0.0001), regardless of *Pfdhfr* haplotype. Parasites with the *Pfdhps* I**<u>AG</u>**KAA haplotype did not have higher SDX EC_50_ values compared to IS**<u>G</u>**KAA parasites. This confirms previous findings that A437G is a key mediator of SDX resistance (51), while S436A is not a mediator of SDX resistance. The new observation in this work is that parasites with the additional A613S mutation (*Pfdhps* I**<u>AG</u>**KA**<u>S</u>**haplotype) had significantly higher EC_50_ values than all other *Pfdhps* haplotypes, suggesting that *Pfdhps* A613S also plays a role in mediating SDX resistance.

#### SP resistance

Given that PYR and SDX are always given in combination, we also wanted to determine the role of the full *Pfdhfr* and *Pfdhps* haplotypes in mediating resistance to SP. Parasites were assayed for SP susceptibility by testing the same concentrations of PYR as was done for the PYR EC_50_ assays, but also adding 100 μM SDX to each of the tested PYR concentrations. The parasite with the highest SP EC_50_ value was *Pfdhfr* **<u>IRN</u>**+ *Pfdhps* I**<u>AG</u>**KAA, suggesting that the accumulation of *Pfdhfr* and *Pfdhps* mutations leads to high SP resistance (figure 4C). Parasites from the same *Pfdhfr* and *Pfdhps* mutation categories (i.e., *Pfdhfr* **<u>IRN</u>** + *Pfdhps* I**<u>AG</u>**KAA, light blue: Th029.10 and Th122.16) had different phenotypes, suggesting that additional differences in the genetic background other than these known mutations (*Pfcrt*, *Pfdhfr*, *Pfdhps*, and *Pfmdr1*) contribute to SP phenotypes. Notably, *Pfdhfr* **<u>IRN</u>** + *Pfdhps* I**<u>AG</u>**KAA parasites did not have statistically significantly different SP EC_50_ values compared to *Pfdhfr* **<u>IRN</u>**+ *Pfdhps* I**<u>AG</u>**KA**<u>S</u>** mutants, whereas the SDX EC_50_ data clearly showed the A613S mutation decreased SDX susceptibility. These finding raise interesting questions about how *Pfdhfr* and *Pfdhps* mutations interact to confer increasing resistance to SP, whether SDX or PYR contributes more to the effectiveness of SP, and how these mutations could affect parasite fitness.

### Pfdhfr *and* Pfdhps *allele population frequency over time*

Given these *in vitro* findings about PYR, SDX, and SP susceptibility, we wanted to investigate the frequency of these different haplotypes in Senegal over time, especially since the introduction of SMC in 2014. The *Pfdhfr* **<u>IRN</u>** mutant haplotype has been close to fixation since 2018 and all other *Pfdhfr* haplotypes have remained at much lower frequencies (supplemental figure 7A). Unlike the *Pfmdr1* and *Pfcrt* combined haplotypes, the *Pfdhfr* and *Pfdhps* combined haplotypes differed between Thiès and Kédougou, possibly due to the different number of monthly SMC rounds at each site (Thiès is a low transmission area with no SMC, while Kédougou is a high transmission area with 5 rounds of monthly SMC) (figure 5, Senegal-wide and Thiès and Kédougou datasets can be found in supplemental figure 7). In Thiès, the most common haplotype is **<u>IRN</u>** + I**<u>A</u>**AKAA (yellow), which has been increasing in frequency since 2012 (supplemental figure 7) and replaced **<u>IRN</u>** + IS**<u>G</u>**KAA (orange) as the most common haplotype in 2020 (figure 5A). In Kédougou, **<u>IRN</u>** + IS**<u>G</u>**KAA (orange) has been the most common haplotype since data began being collected in 2019 and has increased in frequency from 2019 to 2022 while the **<u>IRN</u>** + I**<u>A</u>**AKAA (yellow) haplotype has decreased in frequency (figure 5B). Given the continued use of SP since 2014, it is not surprising to see certain *Pfdhfr* and *Pfdhps* combined haplotyped increasing in frequency in Senegal, especially given our *in vitro* finding that the **<u>IRN</u>** + IS**<u>G</u>**KAA haplotype results in a marked increase in SDX EC_50_ compared to wild-type and the I**<u>A</u>**AKAA haplotype.

**Figure 5.**
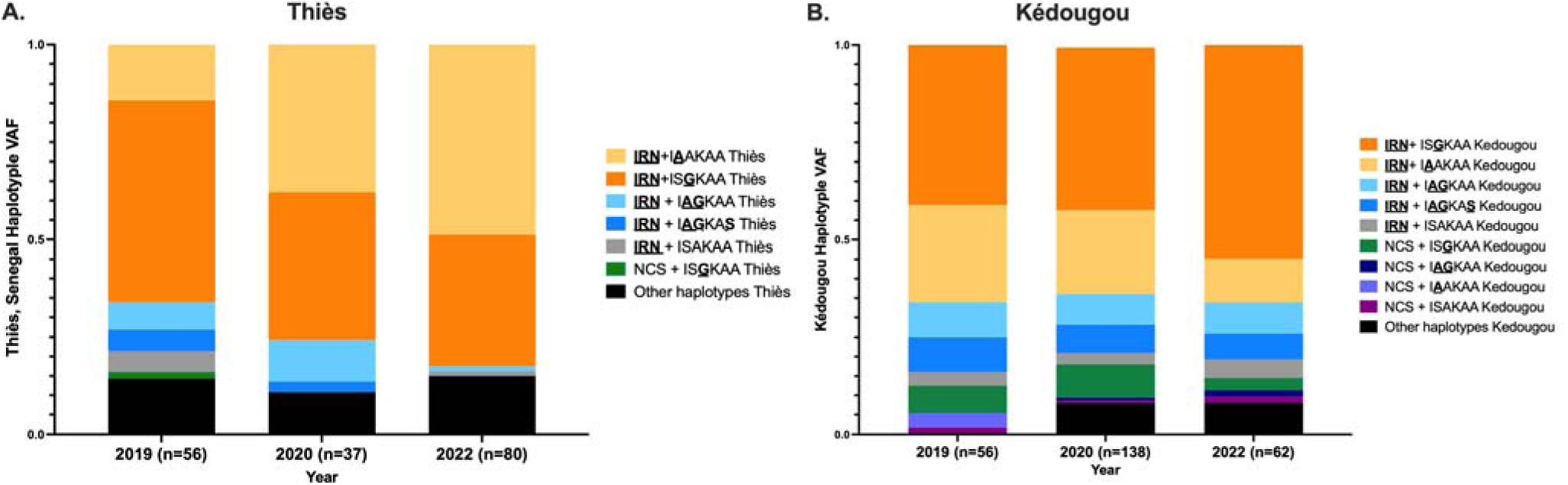
Haplotype variant allele frequencies (VAF) for A. *Pfdhfr* + *Pfdhps* for Thiès and B. *Pfdhfr + Pfdhps* for Kédougou. Samples collected from patients in Thiès and Kédougou from 2019-2022 were whole genome sequenced, monogenomic infections with genotype calls at *Pfdhfr* and *Pfdhps* were included in this dataset. (A) *Pfdhfr* **<u>IRN</u>**+ *Pfdhps* I**<u>A</u>**AKAA is the most prevalent haplotype in Thiès while (B) **<u>IRN</u>** + IS**<u>G</u>**KAA is the most prevalent haplotype in Kédougou. The full list of haplotypes that are included in the “other haplotypes” category can be found in supplemental table 4.

However, what is surprising is that the **<u>IRN</u>** + I**<u>A</u>**AKAA haplotype, which does not confer any additional *in vitro* SDX resistance compared to wild-type *Pfdhps* (ISAKAA), is increasing in frequency. Our analysis of 10,183 samples from the MalariaGEN Pf7 dataset revealed that these major haplotypes found in Senegal are also commonly found in West and Central Africa, however they are not common in East and Northeast Africa and are rarely found in Asia or South America (supplemental figure 8).

### *The role of* Pfgch1 *in pyrimethamine resistance*

*Pfgch1* copy-number amplifications have been shown to decrease susceptibility to PYR depending on the number of *Pfdhfr* mutations (31). In our dataset, we found 3 parasites with *Pfdhfr* **<u>IRN</u>** and *Pfgch1* promoter copy-number amplifications (Th146.12, Th080.12, and Th015.14) and 1 parasite with *Pfdhfr* wild-type and *Pfgch1* gene copy-number amplifications (Th052.16). There was no statistically significant PYR EC_50_ difference between parasites that were *Pfdhfr* **<u>IRN</u>**and had *Pfgch1* promoter copy-number amplifications and parasites that were *Pfdhfr* **<u>IRN</u>** (supplemental figure 9). Therefore, in our dataset, *Pfgch1* copy-number did not explain the range of PYR EC_50_ phenotypes amongst the *Pfdhfr* **<u>IRN</u>** mutant parasites.

### *The* in vitro *fitness* of Pfdhfr and Pfdhps *alleles in the absence of drug*

To investigate the fitness costs of SP resistance, a set of 14 parasites with various *Pfdhfr-Pfdhps* haplotypes (all *Pfcrt* wild-type, CVMNK, given the known fitness costs of *Pfcrt* mutations) were competed in pairwise competitive growth assays for a total of 91 competitions, using the same methods described for the 11 *Pfcrt* CVIET mutants, resulting in a clear ranking (percentage wins) for all 14 parasites. *Pfmdr1* NYD wild-type parasites and *Pfmdr1* NFD mutant parasites both displayed a range of fitness phenotypes (supplemental figure 10), as was seen for the 11 *Pfcrt* CVIET mutant parasites. There was no difference between the relative fitness of *Pfdhfr* NCS wild-type parasites and *Pfdhfr* **<u>IRN</u>** mutants; both genotypes displayed a range of fit and unfit parasites (figure 6A). When considering the relative fitness of individual *Pfdhps* mutations and combinations of *Pfdhps* mutations, there was not a clear pattern. It seems that *Pfmdr1* and *Pfdhfr* play a role in fitness (figure 6B, open shapes are *Pfmdr1* wild-type, triangles are *Pfdhfr* wild-type), as well as components of each parasite’s unique genetic background, making it difficult to ascertain the contribution of *Pfdhps* mutations to fitness. Each genotype examined (whether individual *Pfdhps* mutations, double, or triple) resulted in a range of fitness phenotypes.

**Figure 6.**
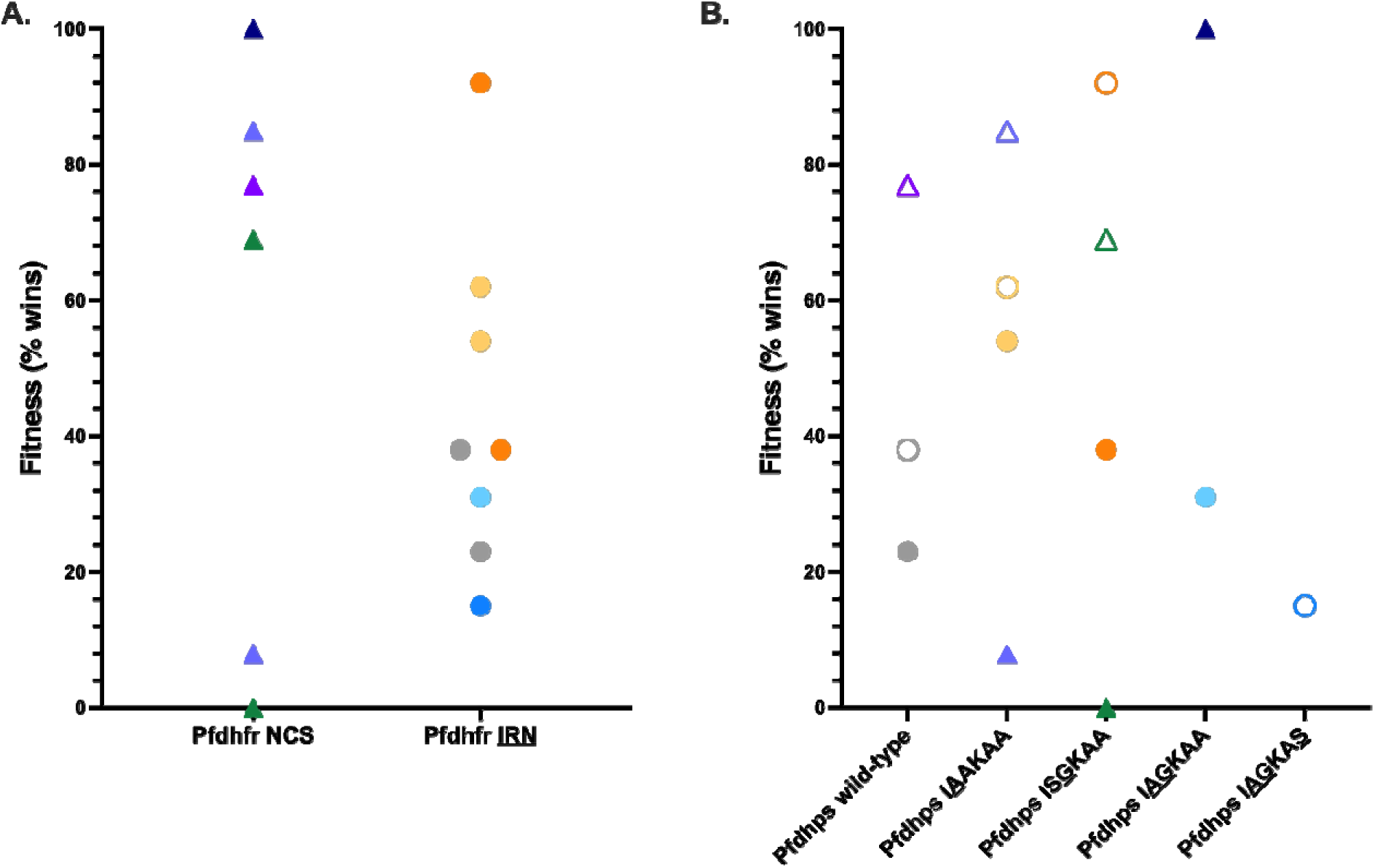
Relative fitness of 14 *Pfcrt* CVMNK parasites based on A. *Pfdhfr* haplotype and B. *Pfmdr1, Pfdhr,* and *Pfdhps* haplotype. (A) Parasites with *Pfdhfr* IRN mutations and *Pfdhfr* wild-type (NCS) parasites both display a range of fitness. Triangles are *Pfdhfr* NCS wild-type. (B) Amongst wild-type *Pfcrt* parasites, a *Pfdhps* I**<u>AG</u>**KAA mutant is the most fit. The least fit parasite had the *Pfdhps* IS**<u>G</u>**KAA genotype. Triangles show *Pfdhfr* NCS wild-type and open shapes show *Pfmdr1* NYD wild-type.

## DISCUSSION

In our study, we found that CVIET + A220S + Q271E + N326S + R371I (FCB) and *Pfcrt* CVIET + A220S + Q271E + I356T + R371I (Cam783) mutants were significantly more resistant to md-AQ compared to *Pfcrt* wild-type and *Pfcrt* CVIET + A220S + Q271E + R371I (GB4) mutants. *Pfdhfr* triple mutants were significantly more PYR resistant than *Pfdhfr* wild-type and revealed a range of resistance phenotypes. *Pfdhps* A437G parasites were significantly more SDX resistant compared to *Pfdhps* wild-type and *Pfdhps* S436A mutants, suggesting that A437G is a key mutation for SDX resistance.

The *Pfdhps* triple mutant (S436A + A437G + A613S) mutation resulted in the highest SDX EC_50_ values. Using competitive growth assays, we found that *Pfcrt* was the main determinant of parasite fitness; there was not a clear correlation between fitness and the other drug resistance markers.

This study contributes to our understanding of drug resistance, which is an evolutionary balance between resistance-conferring mutations and the fitness costs of those mutations. The acquisition of drug-resistance mutations oftentimes imposes a fitness cost to the parasites, which are advantageous when parasites are under drug, but are disadvantageous when parasites are in drug-free conditions (52–54). There have been many examples of drug sensitive parasites returning to high frequency in the population after the discontinuation of the antimalarial drug that selected drug resistant parasites. One such example is the loss of *Pfcrt* mutants in places in the world that replaced CQ as the frontline antimalarial (55–57), and *in vitro* assays have clearly established the fitness cost of *Pfcrt* mutations (32–34). So why do some countries, like Senegal, still have a relatively large frequency of *Pfcrt* mutants (0.49 frequency in 2020 (19))? We hypothesize that these *Pfcrt* mutants remain in the population because they are under selection from amodiaquine use in the population.

Our study presents a unique methodology to answer these questions: we have decades of molecular surveillance data that we can use to inform our hypotheses about what is happening in natural parasite isolates in the population and we can test these hypotheses using *in vitro* culture-adapted natural parasite isolates. Importantly, these natural parasite isolates serve as a “natural experiment”, they are a sample of the viable and successful combinations of genotypes that are circulating in the population. Our set of 33 cryopreserved natural parasite isolates represented 64% of all haplotypes present in our Senegal dataset (supplemental figure 1), thus capturing the genotypic diversity in Senegal and reflecting the consequence of decades of drug pressure in Senegal. This set of parasites allowed us to directly test our hypotheses from molecular surveillance data with *in vitro* phenotypes, as a type of “phenotypic surveillance”.

### Amodiaquine

Previously, AQ resistance has been associated with mutations in *Pfcrt* (K76T) and *Pfmdr1* (N86Y and D1246Y) (9–12). However, using our set of natural parasite isolates, we found that mutant *Pfcrt* CVIET haplotype parasites (regardless of *Pfmdr1* mutation) had increased md-AQ EC_50_ values compared to wild-type *Pfcrt* CVMNK parasites, especially parasites with the Cam783 haplotype (figure 1B), they were relatively fit compared to other *Pfcrt* CVIET parasites (figure 3B), and these parasites have been increasing in frequency in Senegal since 2012 (figure 2, supplemental figure 5).

Parasites with the *Pfcrt* FCB haplotype had the second highest md-AQ EC_50_ values (figure 1B) and the second highest relative fitness rating amongst *Pfcrt* CVIET parasites (figure 3B), however, they remain at low frequencies in Senegal. A previous study using isogenic lines has shown that parasites with the Cam783 haplotype had elevated md-AQ EC_50_ values, but Dd2 and parasites with the FCB haplotype result in the highest md-AQ EC_50_ values, suggesting that the N326S mutation is an important contributor to AQ resistance (32). Interestingly, several studies have found that parasites with the *Pfcrt* FCB haplotype are less fit than parasites with either the *Pfcrt* Cam783 haplotype or parasites with the GB4 haplotype, suggesting that the N326S mutation comes with a fitness cost while the I356T mutation comes with a fitness advantage (32,34), revealing interesting evolutionary trajectories for *Pfcrt* and AQ resistance (33,58). The FCB, GB4, and Cam783 haplotypes have been found to not mediate PIP resistance, but the specific *Pfcrt* mutations in each of these haplotypes are required for further *Pfcrt* mutations to confer PIP resistance (44). Furthermore, the *Pfcrt* N326S and I356T mutations were identified as being part of a genetic background associated with emerging artemisinin resistance (17). Given the continued use of AQ as a component of frontline antimalarials and chemoprevention and given that the major *Pfcrt* mutant haplotypes in Africa are GB4 and Cam783 (supplemental figure 6), it will be important to continue to monitor these *Pfcrt* haplotypes for additional mutations that could change their drug resistance profiles or parasite fitness.

### Sulfadoxine-pyrimethamine

Despite the discontinuation of SP as a first-line therapy in the early 2000s due to the emergence of SP resistance, drug sensitive parasites have not returned to Senegal. This suggests that the use of SP as a chemopreventive is enough to maintain selection for SP resistance; in 2021, almost 45 million children received at least one dose of SMC and nearly 180 million doses were delivered (59). SP is currently used in several African countries in IPTp and SMC, and given the efficacy of SP, the lack of adverse effects, and a lack of suitable drugs ready to replace it, SP will likely continue to be used for years to come (60). SP efficacy is not evaluated through conventional Therapeutic Efficacy Studies (TES), therefore, studies that evaluate the contribution of mutations to resistance and parasite fitness are critical to better understand the role of mutations on parasite survival. Our study assessed the contributions of *Pfdhfr*, *Pfdhps*, and *Pfgch1* to PYR, SDX, and SP resistance. By testing the susceptibility of different *Pfdhfr* + *Pfdhps* haplotypes to PYR, we confirmed that *Pfdhfr* triple mutations drive resistance to PYR, although we found a wide range of phenotypes, suggesting that more than just *Pfdhfr* is playing a role in determining susceptibility to PYR (figure 4A), but it is not *Pfgch1* in our set of parasites (supplemental figure 9). To our knowledge, our study was the first to test the susceptibility of different combinations of *Pfdhfr* + *Pfdhps* haplotypes to SDX. We found that *Pfdhps* S436A does not result in increased SDX resistance, regardless of *Pfdhfr* mutations. However, parasites with *Pfdhps* A437G (regardless of *Pfdhfr* mutations) showed a marked increase in SDX EC_50_ compared to wild-type and *Pfdhps* S436A mutants. Parasites with *Pfdhps* S436A + A437G showed similar SDX EC_50_ values compared to parasites with *Pfdhps* A437G, which further suggested that the S436A mutation does not play a role in SDX resistance in our parasite panel. Interestingly, parasites with *Pfdhps* S436A + A437G + A613S mutations showed the highest SDX EC_50_ values, suggesting that A613S also plays a role in SDX resistance, however EC_50_ values of the *Pfdhps* S436A + A437G + A613S mutants were only 2.1-fold higher than *Pfdhps* A437G mutants (figure 4B).

Our data from samples collected throughout Senegal shows that the *Pfdhfr* triple mutation is nearly fixed, the frequency of the *Pfdhps* IS**<u>G</u>**KAA and I**<u>A</u>**AKAA haplotypes are increasing in Kédougou and Thiès, respectively, and we are beginning to see more parasites with multiple *Pfdhps* mutations (figure 5). A previous study reported an increase in the prevalence of *Pfdhfr* triple mutant parasites with the *Pfdhps* IS**<u>G</u>**KAA haplotype or with the double mutant *Pfdhps* I**<u>AG</u>**KAA haplotype after SMC implementation in Burkina Faso, suggesting that similar trends are happening in other countries (61). These mutations in *Pfdhps* are located in the para-aminobenzoic acid (PABA)-binding pocket, with the A437G mutation resulting in an increased affinity for PABA and a decreased binding affinity for sulfa inhibitors as well as increased enzyme activity (62,63), suggesting that the A437G resistance mutation does not result in a fitness cost for the parasite and could explain why the A437G mutation seems to be a key mutation in *Pfdhps* for further SDX resistance evolution. Our dataset included parasites with *Pfdhps* mutations but no *Pfdhfr* mutations, contrary to what may evolve most commonly. Since *Pfdhfr* is nearly fixed in Senegal, *Pfdhfr* wild-type parasites are rare, but we were able to culture-adapt several of these parasites with and without *Pfcrt* and *Pfdhps* mutations, suggesting that these parasites are in the population and viable. These parasites were fit (especially those that were *Pfmdr1* NYD wild-type) and those that had *Pfdhps* mutations showed a decreased susceptibility for SDX, however not at the same level as their *Pfdhfr* triple mutant counterparts (figure 4B). Despite these natural isolates going against what would be expected evolutionarily, they provided important information about how *Pfdhfr* mutations may also impact SDX resistance.

To further investigate how *Pfdhfr* + *Pfdhps* haplotypes contribute to SP resistance, we assayed parasites to varying doses of PYR with a constant 100μM of SDX. Overall, we saw that parasites with more *Pfdhfr* + *Pfdhps* mutations had increasing SP EC_50_ values, however it was interesting that not all mutations resulted in big SP EC_50_ value changes (figure 4C). Parasites with *Pfdhfr* **<u>IRN</u>** mutations did result in significantly increased SP EC_50_ values compared to wild-type *Pfdhfr* parasites, but the addition of *Pfdhps* mutations did not always result in significantly different EC_50_ values; *Pfdhfr* **<u>IRN</u>**+ *Pfdhps* A437G mutants had similar SP EC_50_ values to *Pfdhfr* **<u>IRN</u>** + *Pfdhps* S436A + A437G + A613S mutants. This brings up several questions about SP and its efficacy and how other drugs in the combination therapy, like AQ, contribute to the benefits provided by SMC in settings with high SP resistance.

The complexity of the folate pathway, the intricate combinations of resistance mutations, and the influence of the concentration of folates and antifolates in the environment makes it difficult to study individual components, therefore, there have been few studies looking at the fitness costs associated with *Pfdhfr* and *Pfdhps* mutations and they often show conflicting results (as reviewed in (64)). While the selective pressure of using SP for SMC could explain the high prevalence of SP resistant parasites, it is also possible that the use of co-trimoxazole (trimethoprim + sulfamethoxazole; there is known cross-resistance between PYR and trimethoprim (65) and SDX and sulfamethoxazole (66) *in vitro*) or the acquisition of other compensatory mutations could also be playing a role in maintaining the high prevalence of SP resistant parasites. In Ethiopia, SP has not been used as a frontline treatment since 2004 and is not used for IPTp, but the high prevalence of SP mutations persists in the population, suggesting that there may be selective pressure from co-trimoxazole or no fitness costs to these SP resistance mutations (67).

Using competitive growth assays, we assessed the fitness of 14 different *Pfdhfr*-*Pfdhps* haplotypes (all wild-type *Pfcrt* given the fitness cost of *Pfcrt* mutations). Interestingly, we found that the differences in fitness were not driven by *Pfdhfr* and *Pfdhps* haplotypes in the absence of drug, in contrast to mutations in *Pfcrt* where overall fitness is affected by *Pfcrt*. This implies that in the absence of drug, there will be no dramatic effect on the population. Interestingly, these haplotypes are maintained and increase in frequency indicating that the effect of drug selection on a population level is likely to be the major driver in determining the allele frequency at these loci. We plan to test this by comparing *in vitro* fitness in the presence of varying concentrations of therapeutic drug.

While *in vitro* phenotypic surveillance can be a great tool, it also has some limitations; further study is needed to determine the correlations between *in vitro* testing and clinical outcomes of drug treatment. Clinical infections are impacted by the complexity of infection, pharmacokinetics, host immune status, and host nutritional status, especially when studying antifolates which are very sensitive to environmental conditions (68). Notably, *in vitro* susceptibility testing of antifolates requires sub-physiological levels of folate and PABA and therefore may not fully reflect *in vivo* conditions.

This phenotypic surveillance work lays the foundation for future work: having used natural parasite isolates to determine *in vitro* fitness and resistance, the next step is use CRISPR-Cas9 to edit mutations on a single parasite background to create a set of isogenic lines to fully understand which haplotypes result in both increased resistance and fitness. These results will help us understand what the next evolutionary step could be for parasites (*e.g.,* additional mutations in *Pfdhps* such as I431V or A581G (61,69), or whether the East African *Pfdhps* K540E mutations could emerge in Senegal) and how it will impact parasite drug resistance and fitness. SMC remains effective in Senegal, as long as the timing and dosing of SMC maintains protection during the high transmission period (70). The expansion of SMC to areas such as Uganda, where moderate SP resistance mutations are prevalent, has been shown to be efficacious so far, likely because high-level SP resistance mutations (*Pfdhfr* I164L and *Pfdhps* A581G) are rare and AQ continues to be effective (71). However, it remains unknown how each individual component of SMC contributes to its overall efficacy, especially in areas with high SP resistance. Nevertheless, as the use and expansion of SMC continues, it is critical to genotypically, phenotypically (*in vitro*), and clinically (*in vivo*, via chemoprevention efficacy studies (72)) monitor these resistance mutations to determine if the efficacy of SP or AQ will be compromised. In conjunction with genotypic surveillance, phenotypic surveillance of natural parasite isolates can assess the impact of drug pressure, identify evolving genetic determinants of drug resistance, and provide known and new molecular markers for ongoing genomic surveillance to monitor and guide the use of drug-based interventions.

## MATERIALS AND METHODS

### Ethics Statement

Samples were obtained from febrile patients who presented at health facilities for care. Informed consent was obtained from all study participants (or from parents/guardians in the patient was a minor). The study protocol was authorized by the Ministry of Health and Social Action in Senegal (SEN 19/49) and approved by the Institutional Review Board of the Harvard T.H. Chan School of Public Health (IRB protocol 16330).

### Sampling

Parasite samples were collected from treatment seeking patients presenting with fever or history of fever within the past 48 hours in Thiès or Kédougou between 2006 and 2022. All patients with positive tests received free malaria treatment in accordance with the National Health Development Policy in Senegal as recommended by the WHO. In addition to slide preparation and RDT, all consenting patients gave venous blood samples. These blood samples were used for whole genome sequencing and for parasite cryopreservation.

### Whole genome sequencing of patient isolates

Samples were subjected to selective whole genome amplification (sWGA) using Phi29 polymerase followed by magnetic bead clean up and quantification, fragmented, and prepared using NEBNext Ultra II FS DNA library as described in Schaffner *et al*. (23). Completed libraries were shipped to the Broad Institute of MIT and Harvard in Cambridge, MA for Illumina-based short read whole genome sequencing. Variant calling was performed in accordance with the best practices established in the Pf3K project using GATK3.5.0 and *Plasmodium falciparum* 3D7v.3 reference assembly as described in Schaffner *et al*. (23). 946 whole genome sequenced monogenomic (single genome) samples were included in our final dataset and used for further analysis.

### Culture adaptation of parasites

Parasites were culture-adapted by thawing cryopreserved *Plasmodium falciparum* isolates collected from patients that had been mixed with glycerolyte, with approximately 1.67 ml of glycerolyte added to every 1 ml of packed RBCs. Samples were stored overnight at -80°C, and then transferred to liquid nitrogen for long term storage. Cryopreserved samples were thawed from liquid nitrogen storage by placing them into a 37 °C water bath. Immediately upon thawing the sample was transferred to a 50 ml conical tube and volume was measured. For every 1 ml of sample volume, a volume of 0.2 ml of sterile 12% NaCl was added dropwise with gentle swirling and then incubated for 5 min at room temperature. Then, for every 1 ml of original sample volume, a total of 9 ml of sterile 1.6% NaCl solution was added with gentle swirling and incubated at room temperature for 2 min. Finally, for every 1 ml of original sample volume, a total of 9 ml of 0.9% NaCl, 0.2% Dextrose was added to the tube with gentle swirling before the tube was centrifuged (2K, 5 min). After aspirating the pellet, the sample was transferred to tissue culture dishes with O+ fresh human blood. For every 1 ml of original sample an individual tissue culture dish was established by adding 1 ml of 50% hematocrit O+ blood (freshly collected, and no more than seven days from collection) along with 10 ml HEPES buffered RPMI media containing 12.5% AB+ human serum (heat inactivated and pooled). Cultures were placed in modular incubators and gassed with 1%O_2_/5% CO_2_/balance N_2_ gas (10 psi for 150 sec) and incubated with rotation (50 rpm) at 37°C. Cultures were settled for 30 min before changing media to retain the maximal amount of culture. Media was changed and smears made daily to monitor parasite growth. 100 μl of O+ RBCs were added to cultures each week while waiting for parasites to reach ∼1% parasitemia (and a first stock freeze). If parasitemia was still low 2 weeks post-thaw, cultures were split 1:2 to add fresh media and RBCs (both splits were kept) to encourage growth.

∼30 stocks of parasites grown in 12.5% AB+ human serum culture media were made by centrifuging (800 x g for 5 min) 9 ml of a predominantly ring stage culture and adding 0.3 ml AB+ human serum for every 0.2 ml of pellet and adding an equal volume of glycerolyte 57 (e.g., 1.0 ml if a 0.4 ml pellet and 0.6 ml of AB+ serum) and aliquoting into a cryotube (∼0.5 ml per aliquot, results in 3-4 stocks) and storing in liquid nitrogen. Several DNA samples (∼9 ml of parasites at 3% or higher parasitemia and late stages were centrifuged at 800 x g for 5 min) were taken and stored in -20 °C freezer until genomic DNA (gDNA) was extracted using the Qiagen QIAamp DNA Blood Mini Kit for genotypic validation.

Parasites were then transitioned to a 50/50 mix of 12.5% AB+ serum culture media and 0.5% Albumax culture media (RPMI 1640 medium supplemented with 28 mM NaHCO_3_, 25 mM HEPES, 400 μM hypoxanthine, 25 μg/mL gentamicin, and 0.5% Albumax II) for 2-3 life cycles and when growing well were transitioned to only 0.5% Albumax media. ∼30 stocks of parasites grown in 0.5% Albumax media were frozen down along with several DNA samples for genotypic validation. All phenotyping was done with parasites grown in 0.5% Albumax media.

### Maintenance of parasites

Once culture adapted, parasites were cultured by standard methods (73) in RPMI 1640 medium supplemented with 28 mM NaHCO_3_, 25 mM HEPES, 400 μM hypoxanthine, 25 μg/mL gentamicin, and 0.5% Albumax II (Life Technologies, Carlsbad, CA) at 5% hematocrit in fresh human O+ erythrocytes (Interstate Blood Band, Inc., Memphis, TN). Cultures were gassed with 1% O_2_/5% CO_2_/balance N_2_ gas and incubated with rotation (50 rpm) in a 37 °C incubator. Cultures were kept below 3% parasitemia with media changes at least every 48 h.

### Parasite genotypic validation

Parasites were whole genome sequenced when collected from the patient as described above. To confirm parasites had the same genomic regions the patient isolate did prior to culture-adaptation, gDNA collected from each natural parasite isolate after culture-adaptation to 0.5% Albumax media was whole genome sequenced. Sequencing libraries were prepared with Illumina’s Nextera XT Kit and DNA libraries were run on an Illumina NovaSeqX Plus as described in (74). All samples were confirmed to have the same genomic regions of interest pre-culture adaptation and post-culture adaptation.

Parasites were also assayed for copy-number variation for *Pfgch1* and *Pfmdr1* by running a Comparative C_T_ experiment using the Applied Biosystems ViiA 7 Real-time PCR system (Life Technologies). Amplification reactions were done in MicroAmp 384-well plates in 10 μl reactions with PowerUp SYBR Green Master Mix (Applied Biosystems), 150 nM of each forward and reverse primer, and 2.35 ng template. *Pfmdr1* forward (5’-TGCATCTATAAAACGATCAGACAAA-3’) and *Pfmdr1* reverse (5’- TCGTGTGTTCCATGTGACTGT-3’) primers were designed after Price et al. (75); *Pfgch1* forward (5’-AAACACCATCTTTTACCTTTTGAA-3’) and *Pfgch1* reverse (5’- AGCATCGTGCTCTTTAACTCC-3’) primers were designed after Kidgell et al. (29); primers for the endogenous control *Pfβ*-tubulin forward (5’-CGTGCTGGCCCCTTTG-3’) and reverse (5’-TCCTGCACCTGTTTGACCAA-3’) were designed after Ribacke et al. (76). Thermocycler conditions were as follows: Uracil-DNA glycosylase (UDG) activation at 50 °C for 2 min, activation at 95 °C for 10 min, 40 cycles of denaturation at 95 °C for 15 sec, annealing at 55 °C for 30 sec, and a final extension at 60 °C for 1 min. Amplification efficiencies were verified by testing a range of gDNA concentrations of all genes and were sufficiently close enough to obviate the need for a correction factor (supplemental figure 11). Copy-numbers were calculated using the ΔΔC_T_ method, ΔΔCt = (Ct_TE_ − Ct_HE_) − (Ct_TC_ − Ct_HC_), where T is the test gene (either *gch1* or *pfmdr1*), H is the reference gene (β-*tubulin*), E is the experimental sample, and C is the control sample. Relative expression was calculated as 2^−ΔΔCt^. Three biological replicates were run per parasite with 0.5 ng/μl gDNA used in each run, each with technical replicates run in quadruplicate; 3D7 and Dd2 controls were included in each run; calculated copy-numbers were averaged. Copy-numbers were considered increased (>1) when the average of the three biological replicates was above 1.6. Assays were repeated if Ct values were greater than 35. Primers were designed to determine whether parasites had amplifications in the *Pfgch1* promoter: forward (5’- GATTCCATTTATTGCATTCTTG-3’) and reverse (5’-CATTTAATGGACTGGAAATT-3’). 25 μl reactions were set up using Phusion High-Fidelity PCR Master Mix with HF Buffer (New England Biolabs, cat #M0531), 0.4 nM of each forward and reverse primer, and 1 μl template. Thermocycler conditions were 98 °C for 90 sec, 30 cycles of 98 °C for 10 sec, 61.4 °C for 2 min 30 sec, and a final extension of 72 °C for 10 min. PCR products were run on a gel, parasites with *Pfgch1* promoter amplifications had a PCR product size of ∼1.5 kb, parasites without *Pfgch1* promoter amplifications had a PCR product of ∼750 bp (supplemental figure 2A). PCR sequencing of 3D7 and Th015.14 was performed by Plasmidsaurus using Oxford Nanopore technology, confirming that Th015.14 has a triple promoter amplification and 3D7 has no amplification (supplemental figure 2B).

### *In vitro* 72 h drug susceptibility assay by SYBR green staining

*In vitro* drug susceptibility of asexual blood stage parasites was measured using the SYBR Green I-based cell proliferation assay as previously described (77). Twenty-four-point dilution series for several of the antimalarial drugs (chloroquine, monodesethyl-amodiaquine, pyrimethamine, and pyrimethamine + 100 μM sulfadoxine) and 12-point dilution curves for the rest of the drugs (mefloquine, lumefantrine, piperaquine, dihydroartemisinin, quinine, and amodiaquine) were carried out in triplicate and repeated with three biological replicates. Each plate included a kill control (negative control) and a no drug control (positive control). All test compounds were resuspended in dimethyl sulfoxide (except for chloroquine, which was prepared in 0.1% Triton X-100 in water and piperaquine, which was prepared in 0.1% Triton X-100 and 0.5% lactic acid in water) and were dispensed into 384-well plates by an HP D300 Digital Dispenser (Hewlett Packard Palo Alto, CA).

Parasites were synchronized to the ring-stage and grown in the presence of different test compounds in 384-clear-bottom well plates at 1% hematocrit, 1% starting parasitemia, and 40 μl of folic acid- and para-aminobenzoic acid-free culture media (RPMI 1640 medium supplemented with 28 mM NaHCO_3_, 25 mM HEPES, 400 μM hypoxanthine, L-glutamate, 25 μg/mL gentamicin, no para-aminobenzoic acid, no folic acid, and 0.5% Albumax II; Gibco, custom order). Growth at 72 h was measured by SYBR Green I (Lonza, Visp, Switzerland) staining of parasite DNA. Relative fluorescence units were measured at an excitation of 494 nm and emission of 530 nm on a SpectraMax M5 (Molecular Devices Sunnyvale, CA). After background subtraction and normalization of raw fluorescence data, half-maximal effective concentration (EC_50_) values were determined using non-linear regression curve fitting in GraphPad Prism 10 (GraphPad Software Inc., Boston, MA). Average EC_50_ values were compared using an Ordinary one-way ANOVA with Tukey’s multiple comparisons test, with a single pooled variance; significance cut-off was p ≤ 0.05.

### Sulfadoxine drug susceptibility

*In vitro* drug susceptibility of asexual blood stage parasites was measured using the SYBR Green I-based cell proliferation assay as described above, with slight modifications for sulfadoxine. A twenty-four-point dilution series for sulfadoxine (resuspended in dimethyl sulfoxide) was carried out in triplicate and repeated with three biological replicates. Parasites were culture adapted to folic acid and para-aminobenzoic acid free culture media (Gibco, custom order) over several life cycles (at least 3 cycles) and were grown with RBCs that were washed 3x in folic acid- and para-aminobenzoic-free RPMI to remove as many folates from the parasite culture as possible prior to being synchronized to the ring-stage for assays. Parasites were assayed at 1% hematocrit, 1% starting parasitemia, and 40 μl of folic acid- and para-aminobenzoic acid-free culture media in 384-well plates. Parasites adapted to folic acid- and para-aminobenzoic acid-free culture media grew noticeably slower, therefore, after 72 h, parasite stage was checked by Giemsa-stained slides. Growth was measured by SYBR Green I as described above, but only when parasites were in late trophozoite/schizont stage, which was an additional ∼8-12 hours after the usual 72 h.

### TaqMan allelic discrimination real-time quantitative PCR-based competitive growth assays

Competitive growth assays between two parasite lines were performed by co-culturing the two parasites in a 1:1 ratio. Parasites were synchronized to the ring-stage and assays were set up at 1% parasitemia (0.5% parasitemia per parasite) and 5% hematocrit. Competitive growth assays were maintained in 2 ml volumes in 12-well plates maintained in regular RPMI + albumax media and kept in a culture chamber (gassed with 1% O_2_/5% CO_2_/balance N_2_ gas in a 37 °C incubator). Each assay had two biological replicates. Plates were maintained with media changes every life cycle and slides were made to monitor parasitemia; at least 75% of the culture volume was collected every life cycle (every 2 days) to isolate gDNA. Competitions were carried out for at least 15 generations. gDNA was extracted using the QIAamp DNA Blood Mini Kit (Qiagen) and quantified via Qubit Fluorometric Quantification.

The percentage of each parasite in each competition was determined using TaqMan allelic discrimination real-time PCR assays on a ViiA 7 Real-Time PCR system. Previously described TaqMan primers (forward and reverse) and TaqMan fluorescence-labeled minor groove binder probes (FAM or HEX) (78) were used to differentiate between each competitor (supplemental table 7 and 8). A standard curve of mixtures of competitor A and B gDNA in fixed ratios (0:100, 20:80, 40:60, 50:50, 80:20, 100:0, and no-template negative control) was run with each set of samples. qPCR reactions for each sample were run in duplicate. 5μl reactions included 2.5μl 2x TaqMan universal PCR master mix, 0.125μl 20x TaqMan-MGB SNP assay mix primers/probes (forward and reverse), and 2.5μl 0.5 ng/μl gDNA. Amplification and detection of fluorescence were carried out using the genotyping assay mode with cycling conditions as follows: 95°C for 10 min, followed by 40 cycles of 95°C for 15 sec and 56°C for 60 sec. Cycle threshold (Ct), the number of cycles required for the fluorescent signal to cross the threshold (i.e., exceeds background level), for each probe (wild-type allele or mutant allele) for each sample was used to determine the wild-type or mutant allele frequency in each sample; Ct values of known mixtures were used to convert Ct values of unknown samples to percentages of wild-type or mutant alleles in each sample. Mixtures containing over 70% of one parasite were counted as a win for that parasite; mixtures containing 29-69% of one parasite after 15 generations were counted as a “tie” for those parasites. Competition outcomes were robust and transitive (e.g., if A beat B and B beat C, A always beat C), allowing for an unambiguous relative fitness ranking for all parasites from the most fit (100% win) to the least fit (0% win).

### Haplotype analysis

To determine the *Pfcrt* + *Pfmdr1* + *Pfdhfr* + *Pfdhps* combined haplotypes from our Senegal dataset, we used the WGS data to call wild-type (0), mutant (1), heterozygous (2) or missing (−1) at each of the 19 sites: *Pfcrt* (codon positions 74-76, A220S, Q271E, N326S, I356T, and R371I), *Pfmdr1* (N86Y, Y184F, D1246Y), *Pfdhfr* (N51I, C59R, S108N), and *Pfdhps* (I431V, S436A, A437G, K540E, A581G, A613S). Based on these calls, any sample that was missing (−1) or heterozygous (2) at any of the 19 sites was removed from the analysis, resulting in 1879 samples. We then determined that our dataset contained 239 unique haplotypes; to determine the most common haplotypes, we counted how many parasites from our dataset had each of these haplotypes. Of these common haplotypes, we selected 33 parasites for culture-adaptation and phenotyping that represented some of the most common haplotypes (these parasites represent 26 different haplotypes and 63.9% of all parasites in our Senegal dataset).

To determine the *Pfcrt* + *Pfmdr1* combined haplotypes over time across Senegal and in just Thiès and Kédougou, we again used our same dataset where we called wild-type (0), mutant (1), heterozygous (2) or missing (−1) at each of the 8 genomic sites: *Pfcrt* (codon positions 74-76, A220S, Q271E, N326S, I356T, and R371I), *Pfmdr1* (N86Y, Y184F, D1246Y). Any sample that was missing or heterozygous was removed from the analysis (resulting in 2160 samples), we then split the data into years: 2005-2006, 2007-2008, 2009, 2010, 2011, 2012, 2013-2014, 2015-2016, 2017-2018, 2019, 2020, 2021, 2022. This data was graphed to represent all samples from all sites in Senegal. The data was then split by collection site and only samples from Thiès were selected for graphing and then only samples from Kédougou were selected for graphing. Similar methodology as described for determining and graphing the *Pfcrt* + *Pfmdr1* combined haplotypes was used for the *Pfdhfr* + *Pfdhps* combined haplotypes, except the 9 genomic sites examined were *Pfdhfr* (N51I, C59R, S108N) and *Pfdhps* (I431V, S436A, A437G, K540E, A581G, A613S), which resulted in 2269 samples.

To determine the *Pfcrt* + *Pfmdr1* combined haplotypes the MalariaGEN Pf7 dataset (79), we called wild-type (0), mutant (1), heterozygous (2) or missing (−1) at each of the 7 genomic sites: *Pfcrt* (codon positions 74-76, A220S, Q271E, N326S, I356T, and R371I), *Pfmdr1* (Y184F). Any sample that was missing or heterozygous was removed from the analysis (resulting in 10,037 samples), we then grouped samples by region: Central Africa (Democratic Republic of Congo), East Africa (Kenya, Madagascar, Malawi, Mozambique, Tanzania), Northeast Africa (Ethiopia, Sudan, Uganda), West Africa (Benin, Burkina Faso, Cameroon, Côte d’Ivoire, Gabon, Gambia, Ghana, Guinea, Mali, Mauritania, Nigeria, Senegal), West Asia (Bangladesh, India, Myanmar), East Asia (Cambodia, Laos, Vietnam, Thailand), Oceania (Indonesia, Papua New Guinea), and South America (Colombia, Peru, Venezuela). To determine the *Pfdhfr* + *Pfdhps* combined haplotypes the MalariaGEN Pf7 dataset (79), we called wild-type (0), mutant (1), heterozygous (2) or missing (−1) at each of the 9 genomic sites: *Pfdhfr* (N51I, C59R, S108N) and *Pfdhps* (I431V, S436A, A437G, K540E, A581G, A613S). Any sample that was missing or heterozygous was removed from the analysis, and samples were grouped by region as described for the *Pfcrt* + *Pfmdr1* combined haplotypes, resulting in 10,183 samples.

## SUPPORTING INFORMATION

**Supplemental figures 1-11.**

**Supplemental table 1.** Full haplotypes (*Pfmdr1* + *Pfcrt* + *Pfdhfr* + *Pfdhps*) represented in our Senegal dataset: frequencies and haplotypes represented by 33 culture-adapted parasites.

**Supplemental table 2.** Full set of phenotype and genotype data for 33 parasite isolates and reference lines.

**Supplemental table 3.** Full list of *Pfmdr1* + *Pfcrt* haplotypes from our Senegal dataset (Senegalwide, Thiès and Kédougou).

**Supplemental table 4.** Full list of *Pfdhfr* + *Pfdhps* haplotypes from our Senegal dataset (Senegalwide, Thiès and Kédougou).

**Supplemental table 5.** Full list of haplotypes from the Pf7 dataset (*Pfmdr1 + Pfcrt* and *Pfdhfr* + *Pfdhps*).

**Supplemental table 6.** Fitness rankings and genotypes for 10 all-on-all pairwise *Pfcrt* mutant competitions (top) and 14 all-on-all pairwise *Pfcrt* wild-type competitions (bottom).

**Supplemental table 7.** List of barcodes used for head-to-head competitions

**Supplemental table 8.** 24 SNP Barcode information for the 25 parasites competed in pairwise competitive growth assays.

## ACKNOWLEDGMENTS

We would like to thank the patients and their families who participated in these studies and the clinic nurses and staff involved with collecting samples from clinics.

## AUTHOR CONTRIBUTIONS

**Conceptualization:** Katelyn Vendrely Brenneman, Wesley Wong, Dyann Wirth, Sarah Volkman

**Funding Acquisition:** Dyann Wirth, Daouda Ndiaye, Sarah Volkman

**Investigation:** Katelyn Vendrely Brenneman, Wesley Wong, Mamy Yaye Die Ndiaye, Karina Bellavia, Imran Ullah

**Methodology:** Katelyn Vendrely Brenneman, Wesley Wong, Stephen Schaffner, Bassirou Ngom

**Resources:** Amy Gaye, Djiby Sow, Mame Fama Ndiaye, Mariama Toure, Nogaye Gadiaga, Aita Sene, Awa Bineta Deme, Baba Dieye, Mamadou Samb Yade, Khadim Diongue, Younouss Diedhiou, Jules François Gomis, Mouhamadou Ndiaye, Mamadou Alpha Diallo, Ibrahima Mbaye Ndiaye

**Supervision:** Bronwyn MacInnis, Dyann Wirth, Daouda Ndiaye, Sarah Volkman

**Writing – original draft:** Katelyn Vendrely Brenneman, Sarah Volkman

**Writing – review & editing:** All authors

