## Supplemental Figures 1-11 for "Phenotypic assessment and genetic validation of *Plasmodium falciparum* molecular markers associated with malaria chemoprevention in Senegal"

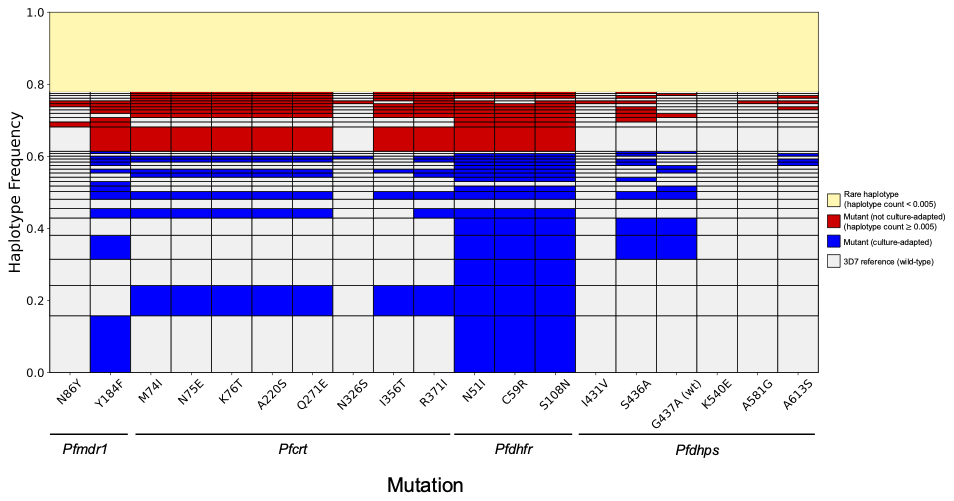

**Supplemental figure 1. Haplotypes represented in our Senegal dataset.** *Pfmdr1* + *Pfcrt* + *Pfdhfr* + *Pfdhps* combined haplotypes that exist in our Senegal dataset from 1879 monogenomic samples. Gray bars represent wild-type alleles at each position, blue or red bars represent mutant alleles at each position. Yellow haplotypes are rare (haplotype count is less than 0.005), haplotypes in red were not culture-adapted (haplotype count is greater than or equal to 0.005), haplotypes in blue include the 33 culture-adapted parasites and represent 26 different haplotypes and 63.9% haplotype frequency.

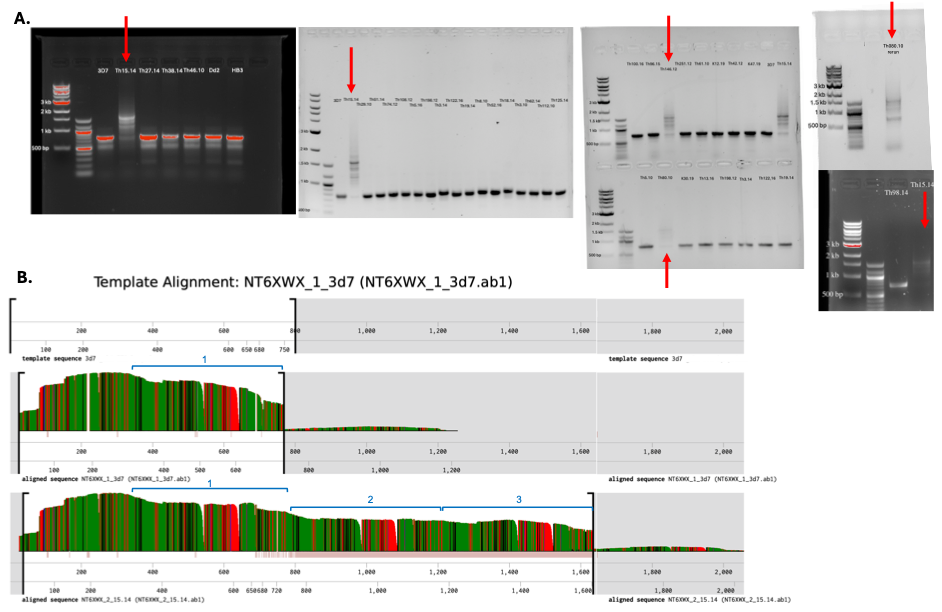

**Supplemental figure 2.** **Evidence of *Pfgch1* promoter amplifications in three parasite isolates.** The promoter region of the *Pfgch1* gene was amplified using PCR and products were run on a gel with an expected size of ~750 bp for a single *Pfgch1* promoter + ~300 bp for every additional copy of the *Pfgch1* promoter present. (A) PCR products from 33 isolates, 3D7, Dd2, and HB3 were run on gels, revealing Th015.14, Th146.12, and Th080.10 to have product sizes around ~1500 bp (red arrows), suggesting a triple promoter amplification in each of these isolates. (B) Custom primers designed to amplify a 772 bp region (Pf3D7_12_v3 973584-974355 (+)) were used to generate PCR products from 3D7 (top) and Th015.14 (bottom). These products were Nanopore sequenced and aligned, revealing that 3D7 indeed has one copy of the *Pfgch1* promoter (resulting in a ~750 bp product), while Th015.14 has three copies of the *Pfgch1* promoter (resulting in a ~1500 product). An approximation of the copied region is marked with blue brackets; 3D7 has one of these regions, while Th015.14 has three repeats of this region. Read depth is depicted as height; regions in gray (far right region) have too few reads and are considered noise.

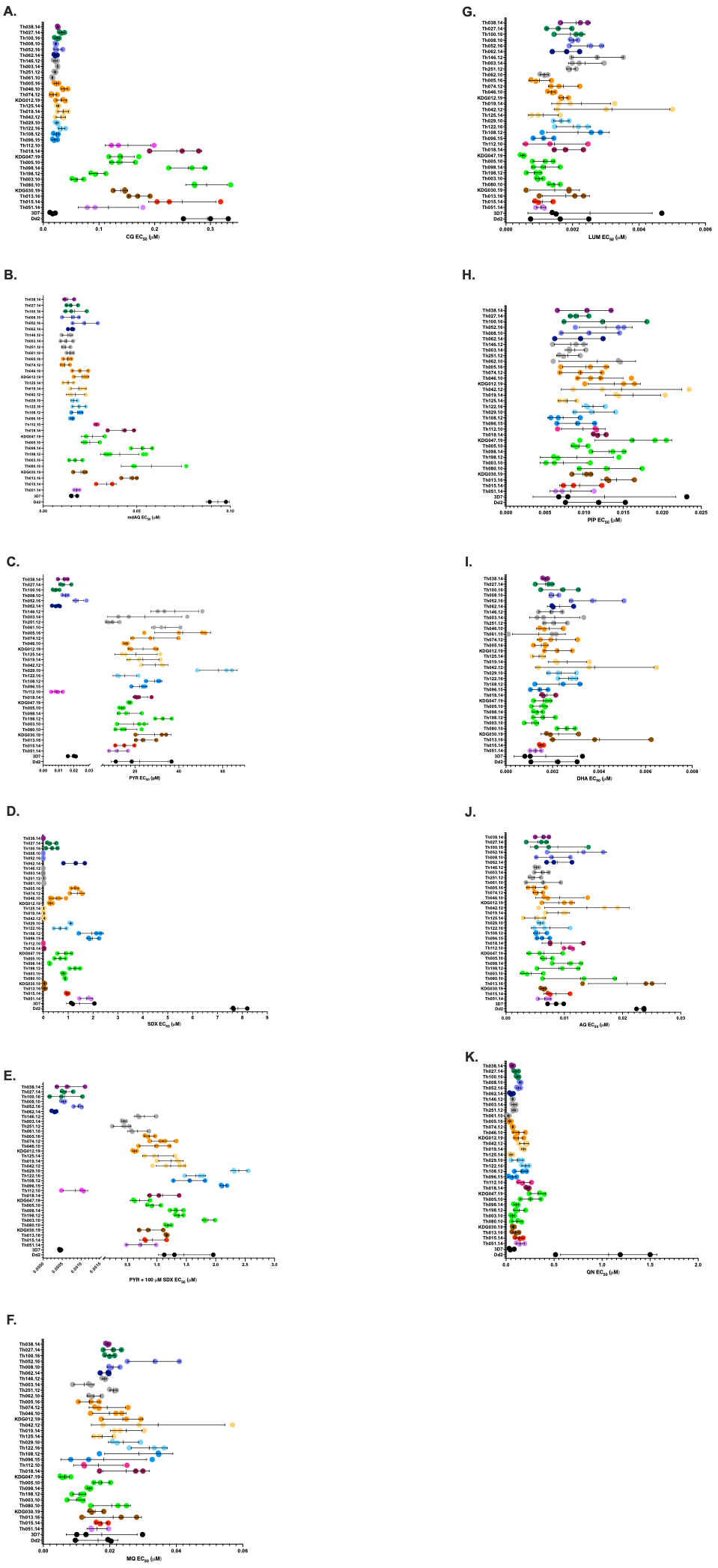

**Supplemental figure 3. EC_50_ values for all parasites and all drugs phenotyped.** EC_50_ values (x-axis) for each biological replicate for 33 culture-adapted parasites and two reference lines (y-axis); genotypes of each parasite are listed in Table 1. For each of the antimalarials assayed, concentration-response curves were generated for at least 3 biological replicates (each with 3 technical replicates) to calculate the EC_50_ value, represented as a dot. Each parasite is shown with the mean and standard deviation of the biological replicates for the following antimalarials: (A) Chloroquine, (B) monodesthyl-amodiaquine, (C) pyrimethamine, (D) sulfadoxine, (E) pyrimethamine + 100 μM sulfadoxine, (F) mefloquine, (G) lumefantrine, (H) piperaquine, (I) dihydroartemisinin, (J) amodiaquine, and (K) quinine. Mean EC_50_ values, standard deviation, and replicates are listed in supplemental table 2. These mean EC_50_ values were used to generate figure 1 and figure 4.

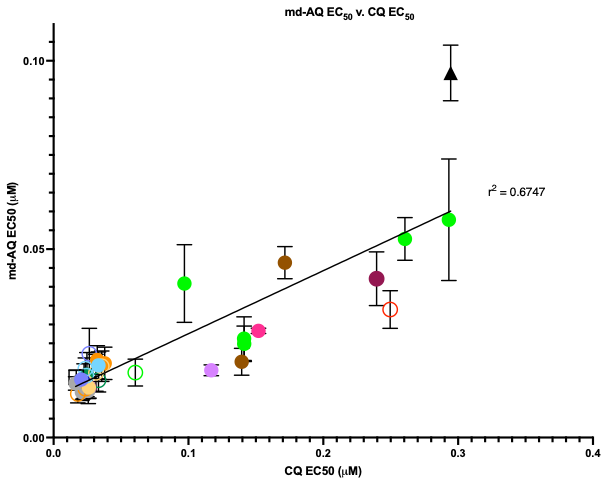

**Supplemental figure 4. Correlation between monodesethyl-amodiaquine (md-AQ) and chloroquine (CQ) EC_50_ values.** md-AQ and CQ EC_50_ values for all 33 culture-adapted parasites and 2 reference lines correlated with a r^2^ value of 0.675, showing that CQ and md-AQ EC_50_ values are positively correlated. Parasites with open circles are *Pfmdr1* NYD wild-type.

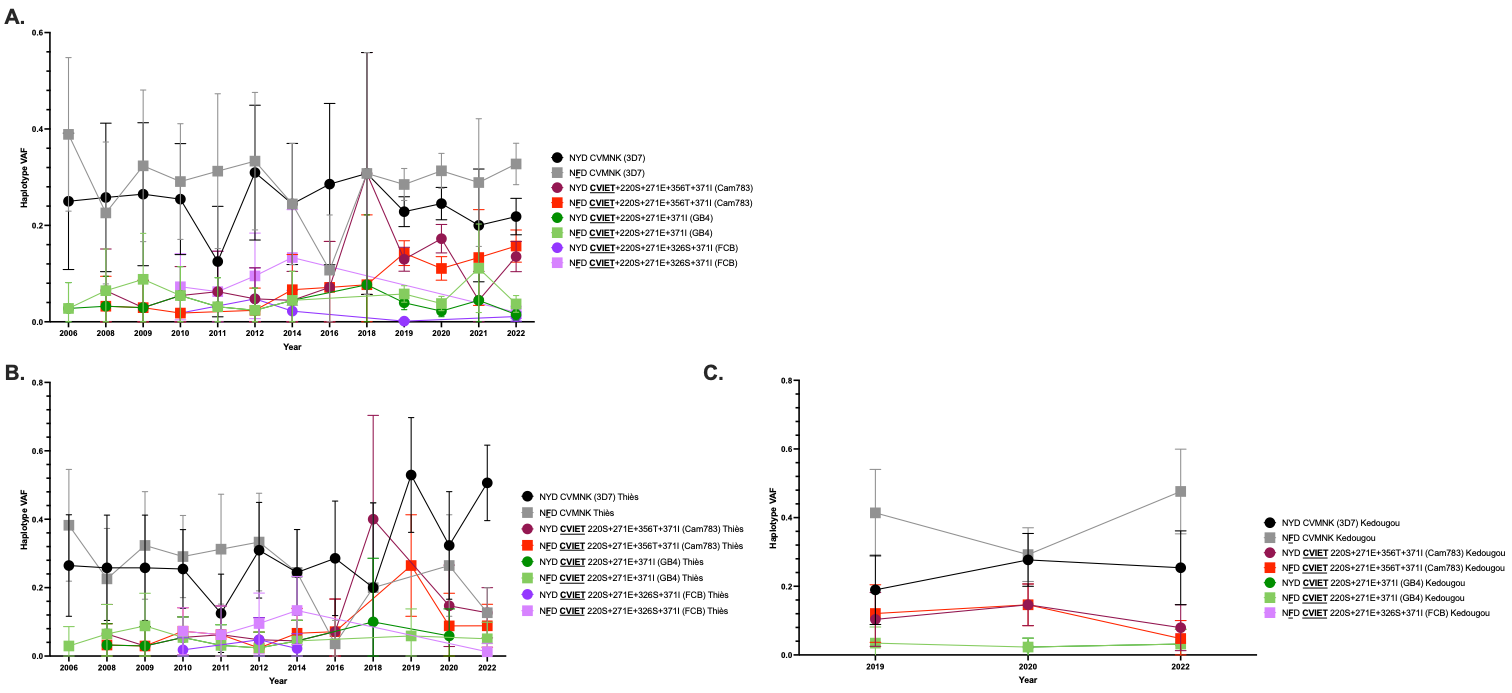

**Supplemental figure 5. *Pfmdr1* + *Pfcrt* combined haplotypes over time from 2160 samples in Senegal.** Each color shows a different haplotype; circles show wild-type *Pfmdr1,* and squares show Y184F mutant *Pfmdr1*. Each year (x-axis) shows the mean haplotype variant allele frequency (VAF) and the 95% confidence interval (y-axis). (A) *Pfmdr1* + *Pfcrt* combined haplotypes frequencies from all 2160 samples from all sites in Senegal, (B) *Pfmdr1* + *Pfcrt* combined haplotypes from samples only from Thiès (458 samples), and (C) *Pfmdr1* + *Pfcrt* combined haplotypes only from samples in Kédougou (251 samples). The full list of haplotypes can be found in supplemental table 3.

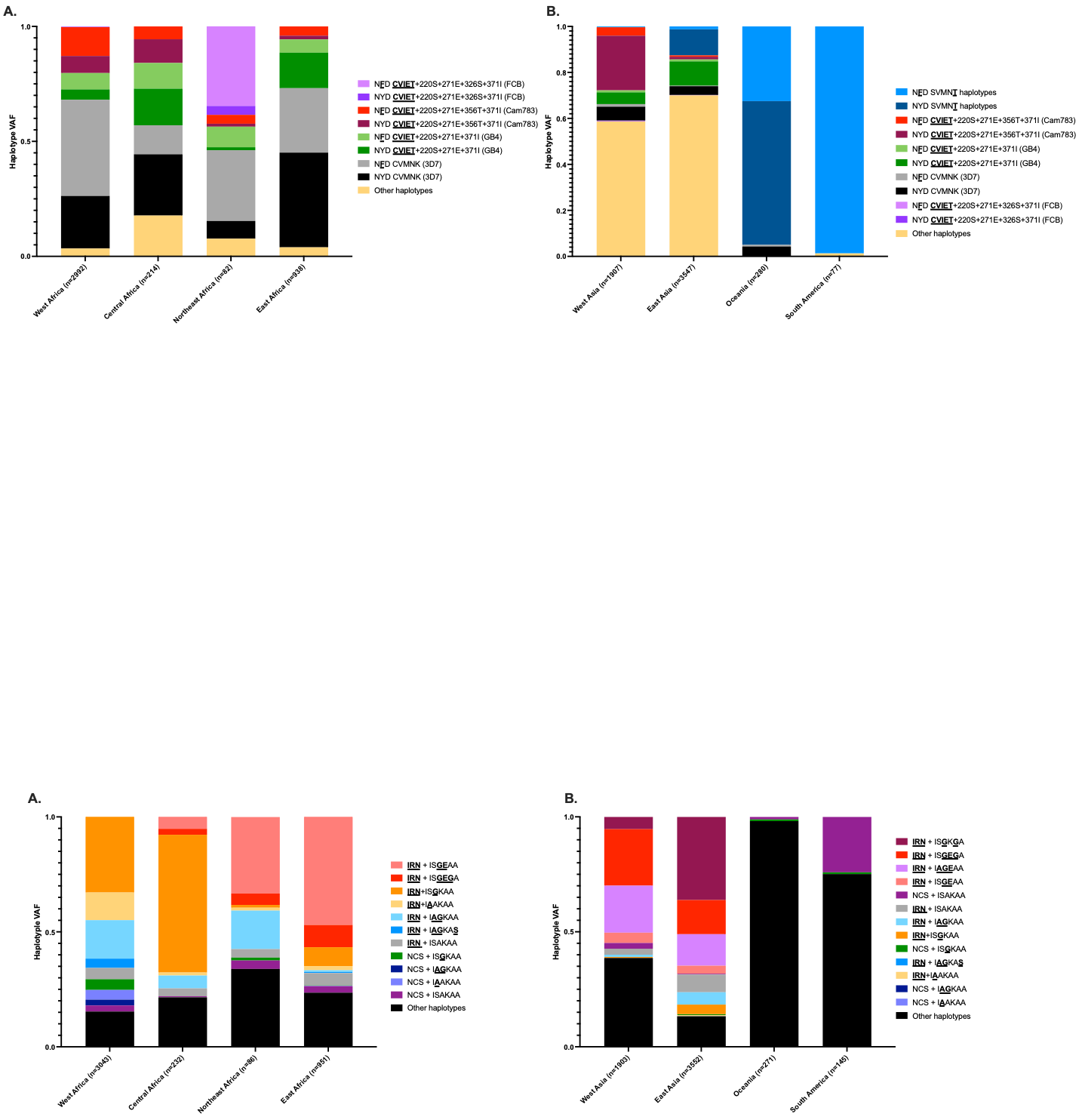

**Supplemental figure 6. *Pfmdr1* + *Pfcrt* combined haplotypes from 10,037 MalariaGEN Pf7 samples.** Graphs show the *Pfmdr1* + *Pfcrt* combined haplotype variant allele frequency (y-axis) for different haplotypes in different regions of the world (x-axis). (A) *Pfmdr1* + *Pfcrt* combined haplotypes for 4 different African regions (West, Central, Northeast, and East; 4226 samples) and (B) *Pfmdr1* + *Pfcrt* combined haplotypes for 4 different regions outside of Africa (West Asia, East Asia, Oceania, South America; 5811 samples). The full list of haplotypes that are included in the other haplotypes category can be found in supplemental table 5.

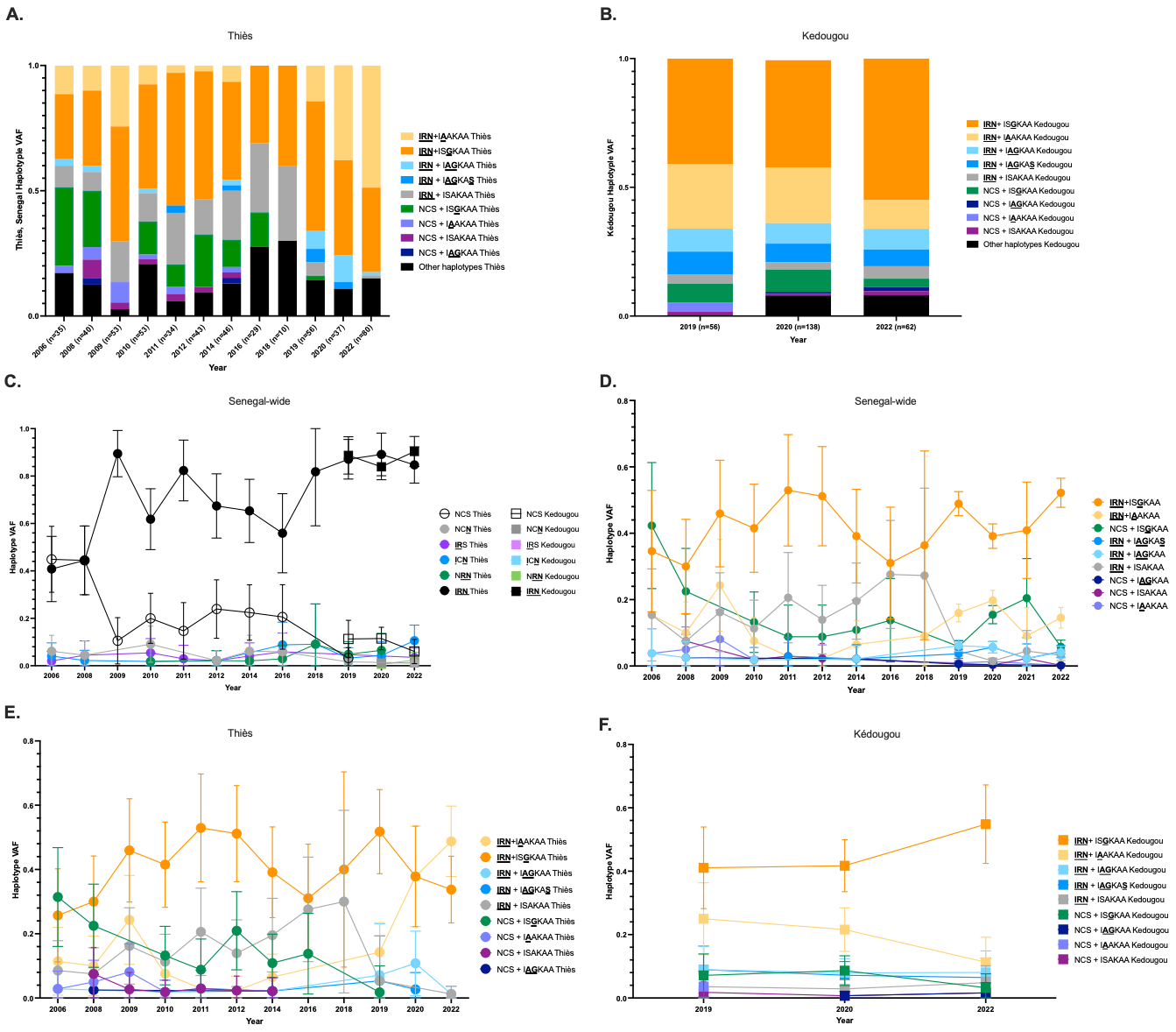

**Supplemental figure 7. *Pfdhfr* + *Pfdhps* combined haplotype frequencies over time in Senegal.** Samples collected from patients in Thiès and Kédougou were whole-genome sequenced, monogenomic infections with genotype calls at *Pfdhfr* and *Pfdhps* were included in this dataset. (A) *Pfdhfr* **IRN** + *Pfdhps* I**A**AKAA is the most prevalent haplotype in Thiès while (B) **IRN** + IS**G**KAA is the most prevalent haplotype in Kédougou. (C-F) Each color shows a different haplotype; circles show samples from Thiès and squares show samples from Kédougou. Each year (x-axis) shows the mean haplotype variant allele frequency (VAF) and the 95% confidence interval (y-axis). (C) *Pfdhfr* haplotype frequencies in Thiès (circles) and Kédougou (squares). (D) *Pfdhfr* + *Pfdhps* combined haplotype frequencies from all 2269 samples from all sites in Senegal, (E) *Pfdhfr* + *Pfdhps* combined haplotypes from samples in Thiès (500 samples), and (F) *Pfdhfr* + *Pfdhps* combined haplotypes from samples in Kédougou (256 samples). The full list of haplotypes can be found in supplemental table 4.

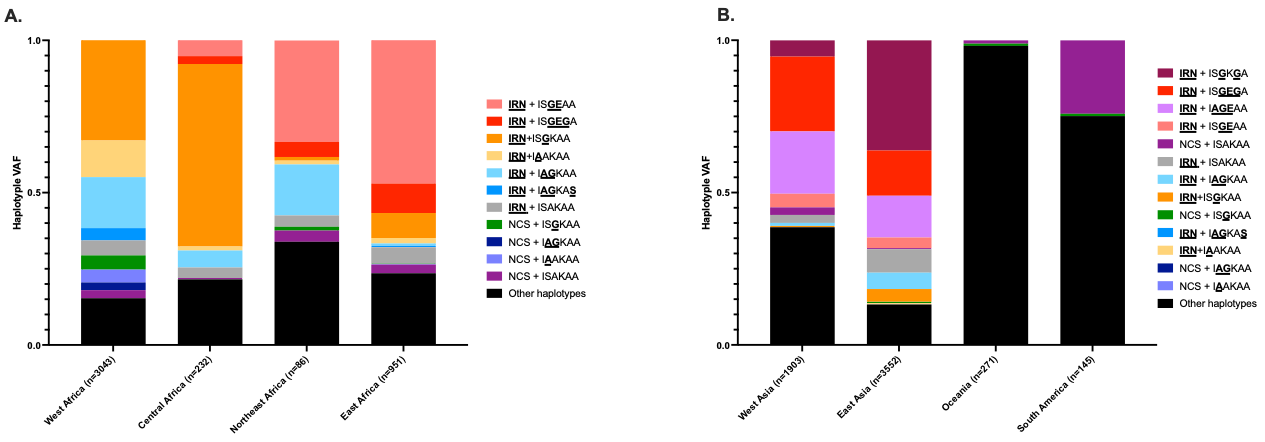

**Supplemental figure 8. *Pfdhfr* + *Pfdhps* combined haplotypes from 10,183 MalariaGEN Pf7 samples.** Graphs show the *Pfdhfr* + *Pfdhps* combined haplotype variant allele frequency (y-axis) for different haplotypes in different regions of the world (x-axis). (A) *Pfdhfr* + *Pfdhps* combined haplotypes for 4 different African regions (West, Central, Northeast, and East; 4312 samples) and (B) *Pfdhfr* + *Pfdhps* combined haplotypes for 4 different regions outside of Africa (West Asia, East Asia, Oceania, South America; 5871 samples). The full list of haplotypes that are included in the other haplotypes category can be found in supplemental table 5.

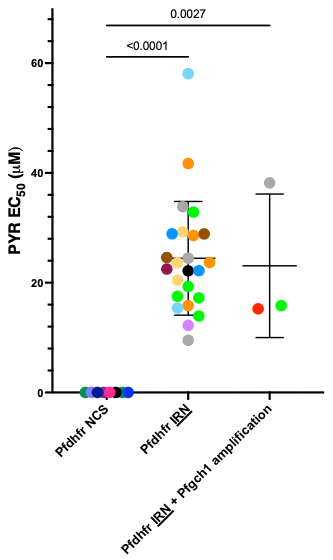

**Supplemental figure 9. Pyrimethamine EC_50_ comparisons between parasites with *Pfdhfr* mutations and *Pfgch1* copy-number amplifications.** Pyrimethamine (PYR) EC_50_ is shown on the y-axis, with different genotypes of parasites on the x-axis. While triple mutant *Pfdhfr* **IRN** parasites were significantly more PYR resistant (p<0.0001), parasites with *Pfgch1* copy-number amplifications did not have a significantly different PYR EC_50_ when compared to parasites with only *Pfdhfr* **IRN** mutations (p=0.0027).

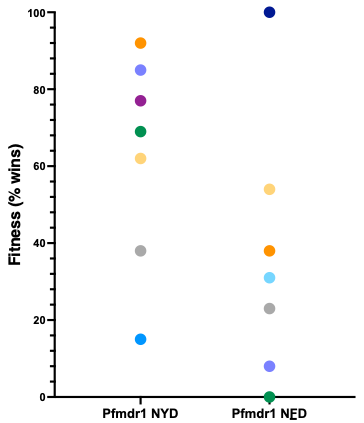

**Supplemental figure 10. Relative fitness ratings of 14 *Pfcrt* wild-type parasites.** Parasites with *Pfcrt* CVMNK (wild-type) and various *Pfdhfr-Pfdhps* haplotypes were competed in all-on-all pairwise competitions to determine each parasite’s relative fitness. Parasites with a higher win percentage (y-axis) were more fit. Parasites with *Pfmdr1* NYD displayed a range of fitness, as did the mutant *Pfmdr1* NFD parasites, suggesting that there is not a particular cost or benefit to either of these genotypes.

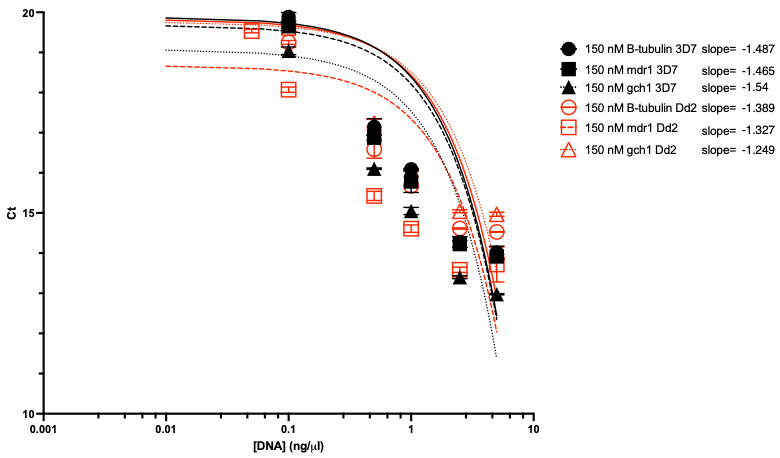

**Supplemental figure 11. qPCR amplification efficiency of *Pfβ-tubulin*, *Pfmdr1*, and *Pfgch1* primers.** Amplification efficiency was verified by testing the *Pfβ-tubulin*, *Pfmdr1*, and *Pfgch1* qPCR primers on a range of DNA concentrations of the reference lines 3D7 and Dd2 (0.01 ng/μl – 2.5 ng/μl). Cycle threshold (Ct; number of cycles of PCR needed to replicate enough DNA to cross the threshold line) value is on the y-axis with the DNA concentration of the sample tested on the x-axis. All primers were tested at 150 nM concentrations. Circles show the amplification of *Pfβ-tubulin* using 3D7 (black) and Dd2 (red), squares show the amplification of *Pfmdr1* using 3D7 (black) and Dd2 (red), and triangles show the amplification of *Pfgch1* using 3D7 (black) and Dd2 (red).
